# CELLector: Genomics Guided Selection of Cancer *in vitro* Models

**DOI:** 10.1101/275032

**Authors:** Hanna Najgebauer, Mi Yang, Hayley E Francies, Clare Pacini, Euan A Stronach, Mathew J Garnett, Julio Saez-Rodriguez, Francesco Iorio

## Abstract

The selection of appropriate cancer models is a key prerequisite for maximising translational potential and clinical relevance of *in-vitro* oncology studies.

We developed *CELLector*: a computational method (implemented in an open source R Shiny application and R package) allowing researchers to select the most relevant cancer cell lines in a patient-genomic guided fashion. CELLector leverages tumour genomics data to identify recurrent sub-types with associated genomic signatures. It then evaluates these signatures in cancer cell lines to rank them and prioritise their selection. This enables users to choose appropriate models for inclusion/exclusion in retrospective analyses and future studies. Moreover this allows bridging data from cancer cell line screens to precisely defined sub-cohorts of primary tumours. Here, we demonstrate usefulness and applicability of our method through example use cases, showing how it can be used to prioritise the development of new in-vitro models and to effectively unveil patient-derived multivariate prognostic and therapeutic markers.

**Graphical Abstract:** 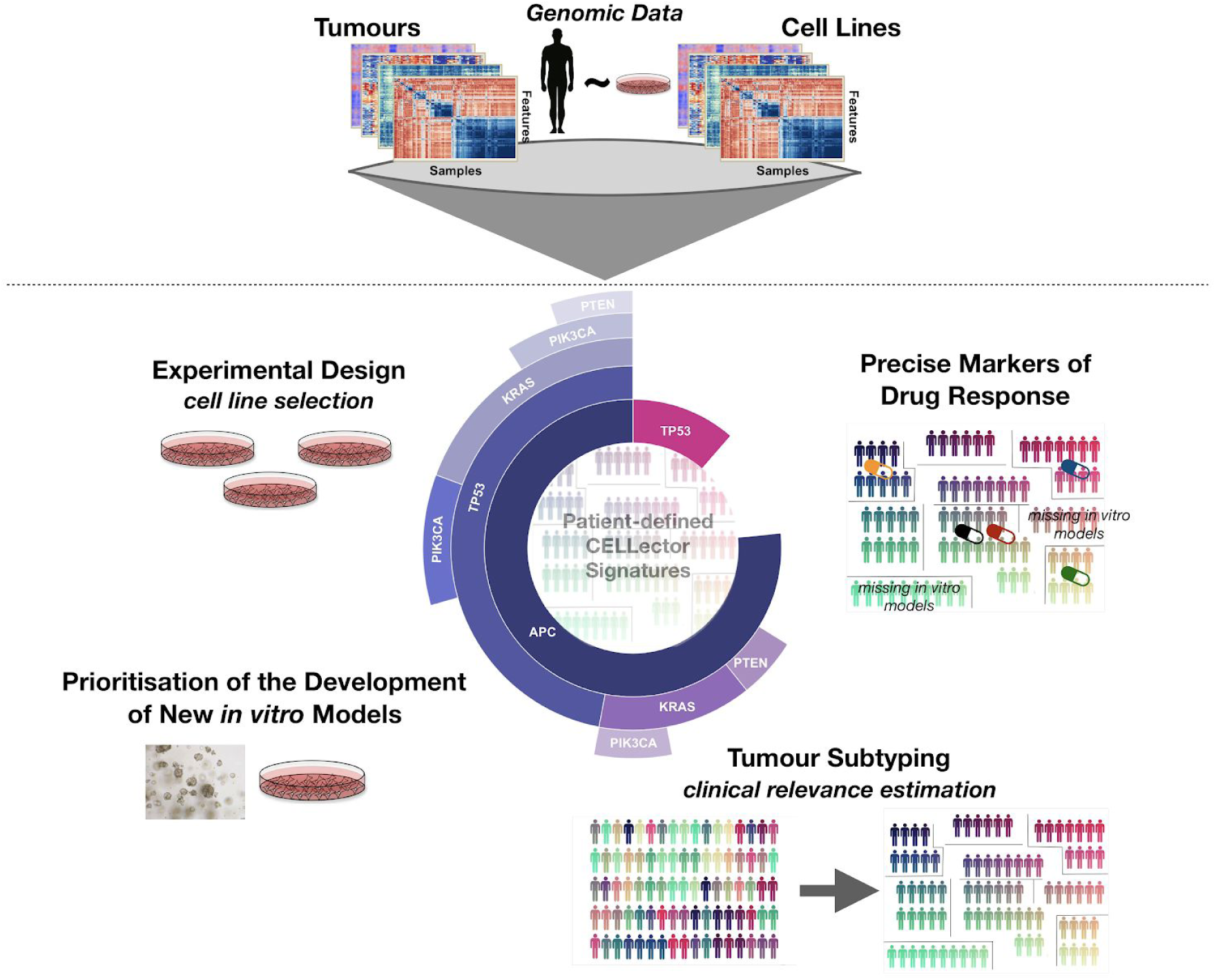

## Introduction

The use of appropriate cancer *in vitro* models is one of the most important requirements for investigating cancer biology and for successfully developing new anticancer therapies. Much effort has been devoted to evaluating the extent of phenotypic and genotypic similarities between existing cancer models and the primary tumours they aim to represent (Ahmed et al., 2013; Beaufort et al., 2014; Ince et al., 2015; Medico et al., 2015; Qiu et al., 2016). Despite inherent limitations, immortalised human cancer cell lines are the most commonly used experimental models in pre-clinical oncology research. Technological advancement in high-throughput ‘omics’ techniques and the availability of rich cancer genomics datasets, such as those provided by The Cancer Genome Atlas (TCGA, http://cancergenome.nih.gov), the International Cancer Genome Consortium (ICGC) (Zhang et al., 2011), the NCI-60 panel (Shoemaker, 2006), the Cancer Cell Line Encyclopedia (CCLE) (Barretina et al., 2012), the Cell Model Passports (van der Meer et al., 2019), the Genomics of Drug Sensitivity in Cancer (GDSC) (Garnett et al., 2012; Iorio et al., 2016), the COSMIC Cell Line Project (Forbes et al., 2017) and many others, have transformed the way preclinical cancer models can be assessed and prioritised. To this end, analytical methods to evaluate the suitability of cell lines as tumour models have been proposed in recently published works (Domcke et al., 2013; Jiang et al., 2016; Mouradov et al., 2014; Sinha et al., 2017; Sun and Liu, 2015; Vincent et al., 2015; Zhao et al., 2017). Although these studies provide useful guidelines for choosing appropriate (and avoiding poorly suited) cell line models, they are restricted to individual cancer types. Most importantly, they require expert knowledge of the genomic alterations known to have a specific functional role in the tumour (sub)type under consideration (Domcke et al., 2013; Jiang et al., 2016), or in the opposite case, they consider all individual variants regardless of their clinical relevance or functional impact, weighting these solely based on their frequency in single-sample based metrics (Sinha et al., 2015). As a consequence, there is a need for robust computational methods able to integrate the molecular characterisation of large cohorts of primary tumours from different tissues, extracting the most clinically relevant features in an unbiased way, and evaluating/selecting representative *in vitro* models on the basis of these features.

We have recently published a large molecular comparison of cancer cell lines and matched primary tumours at the population level (Iorio et al., 2016). Our results show that cell lines recapitulate most of the oncogenic alterations identified in matched primary tumours, and at similar frequencies. Building on our previous work, here we present CELLector, a tool for genomics-guided selection of cancer *in vitro* models.

CELLector is based on an algorithm that combines methods from graph theory and market basket analysis (Han et al., 2011). It makes use of large-scale tumour genomics data to explore and rank patient subtypes based on genomic signatures (e.g. combinations of genomic alterations) identified in an unsupervised way based on their prevalence. Subsequently, it ranks cell line models based on their genomic resemblance to the identified patient subtypes. Additionally, CELLector enables the identification of disease subtypes currently lacking representative *in vitro* models, which could be prioritised for future development. Here we demonstrate clinical relevance and potential translational application of the patient-defined signatures uncovered by CELLector through systematic analyses associating the signatures to differential patient prognosis, and response to *in vitro* drug treatment.

CELLector is available as an open-source user-friendly R Shiny application at https://ot-cellector.shinyapps.io/cellector_app/ (code available at https://github.com/francescojm/CELLector_App) and R package at https://github.com/francescojm/CELLector, it also provides interactive visualisations and intuitive explorations of results and underlying data.

## Results

### Overview of CELLector

CELLector is implemented into two distinct modules. The first module recursively identifies the most frequently occurring sets of molecular alterations (signatures) in a cohort of primary tumours (from TCGA or provided by the user), by focusing on a set of recently published (Iorio et al., 2016) or user-defined clinically relevant genomic features. In the default setting, these genomic features encompass somatic mutations in 470 high-confidence cancer driver genes and copy number gains/losses of 425 recurrently altered chromosomal segments, and were identified by applying state-of-art computational tools, such as the intOGen pipeline (Gonzalez-Perez et al., 2013; Gundem et al., 2010) and ADMIRE (van Dyk et al., 2013), to the genomic characterisation of a cohort of 11,289 cancer patients (from the TCGA (http://cancergenome.nih.gov), the ICGC (Zhang et al., 2011) and other publicly available studies. Epigenomic data can also be used by including in the analysis the (discrete) methylation status of 378 *informative* CpG islands within gene promoters. These were identified in Iorio et al., 2016 based on the multimodal distribution of their methylation signal (indicative of the signal being informative and not tissue-specific) across primary tumour samples in at least one cancer type.

Based on the collective presence/absence of these alterations sets, CELLector partitions the primary tumours into distinct subpopulations prioritising them based on their prevalence in the patient population (Figure 1A). The second module determines the status of the identified molecular signatures (i.e. combinations of genomic alterations) in cancer cell lines (from user-defined or derived from Iorio et al., 2016 data) in order to identify the best-representative models for each patient subpopulation (Figure 1B). This approach unveils the extent of disease heterogeneity covered by representative models, and it also enables the identification of molecular signatures underlying tumour subtypes currently lacking representative models (Figure 1C).

**Figure 1.**
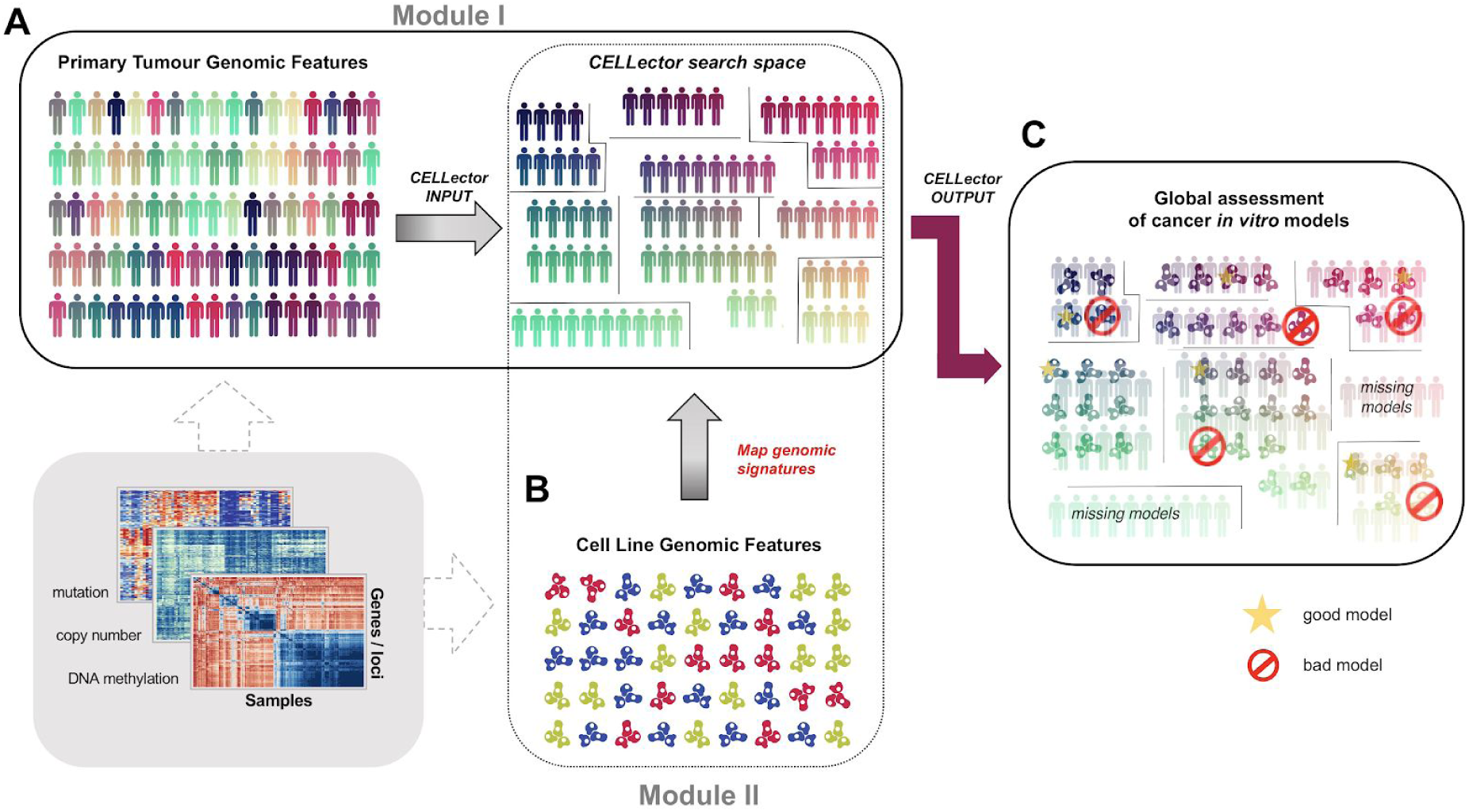
Schematic representation of the CELLector modules. A. Genomic features of primary tumours are used to identify recurrent subpopulations associating them with combination of genomics events (CELLector signatures). These are modeled and visualised hierarchically, by assembling the CELLector search space. B. The resulting CELLector search space is then used to map molecular similarities between the identified tumour subpopulations and cell line models. C. CELLector returns a list of cell line models that best represent the identified tumour subpopulations, thus maximising the coverage of disease heterogeneity, and highlighting tumour subtypes currently lacking representative *in vitro* models.

By default, CELLector makes use of built-in datasets from the genomic characterisation of primary tumours and cell lines derived from 16 different tissues (Supplemental Information Table S1 and STAR Methods). In addition, CELLector can be used on user-defined/provided tumour and cell line genomic datasets, and sets of genomic features and individual genomic variants passing recurrence based filters (based on frequencies observed in COSMIC (Forbes et al., 2017), or user defined ones). Finally, cell line annotations and genomic data can be optionally synchronised to the latest installment of the Cell Model Passports (van der Meer et al., 2019), and an independent module (the CELLector Binary Event Matrix (BEM) builder module) allows creation of fully customisable genomic matrices using public or user-defined genomic data for both primary tumours and cell lines.

#### CELLector modules

In the first module, CELLector assembles a search space by segmenting cancer patients into a hierarchical structure visualised through a *sunburst* diagram. By mapping cell lines onto individual patient segments, this structure provides a prioritisation strategy for choosing the most representative models to be included in a new *in vitro* study. This strategy also ensures that the number of patients represented by at least one selected model, is maximised.

Particularly, CELLector recursively segments patients by grouping them according to the presence/absence of genomic alterations most frequently occurring in the patient cohort. This also minimises the corresponding number of segments, and is performed by a greedy algorithm that proceeds as follows. Starting from an initial cohort of patients, the most frequent genomic alteration (or set of genomic alterations) is identified. This can be an individual mutation/copy-number-alteration/hypermethylated-gene-promoter, or a pair/triplet of such alterations occurring simultaneously. In the latter case these are identified by using the *Eclat* algorithm (Kaur, 2014) for the identification of frequent itemsets in commercial transaction databases. Based on this, the cohort of patients is split into two subpopulations depending on the presence/absence of the most frequent alteration identified (collective presence/absence if considering pairs/triplets of alterations). The obtained partition defines two subsets: the alteration(s) *support* set and its *complement*. The algorithm is then executed recursively on the two resulting subpopulations: to refine the support set (*refinement recursion*) and to analyse its complement (*complement recursion*), respectively. The recursions continue until all the alteration sets with a support of minimal user-defined size are identified, and the corresponding patient segments are defined. The identified patient sub-populations and underlying signatures are stored in a hierarchical data object, which can be visualised as a sunburst and whose structure reflects the recursive calls of the algorithm (Figure 2A).

**Figure 2.**
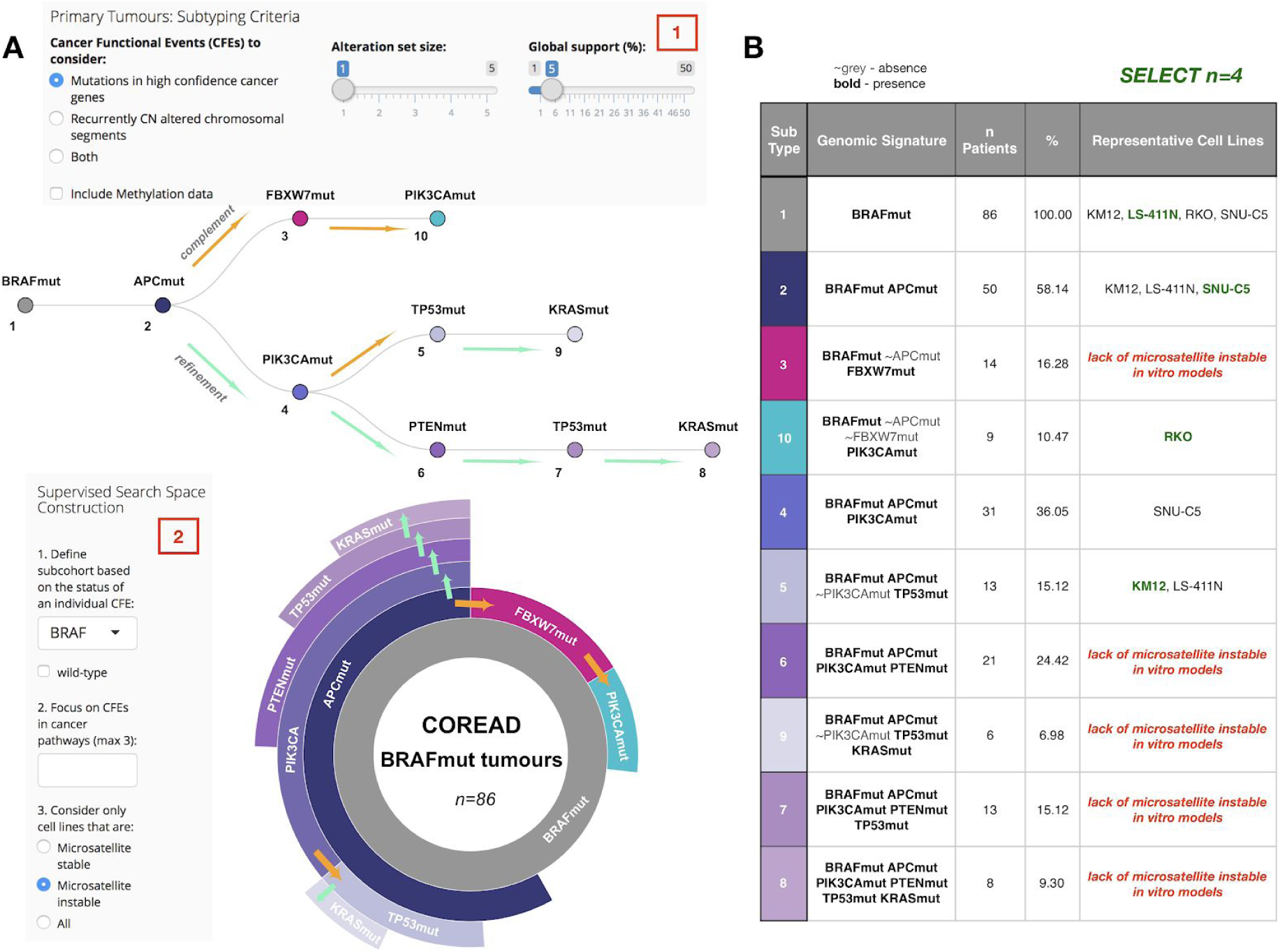
CELLector interface with an example case study. **A.** Visual representation of the CELLector search space constructed based on the prevalence of mutations in high-confidence cancer driver genes observed in cohort of *BRAF* mutant colorectal (COREAD) tumours (box 1 and box 2), and considering only microsatellite instable cell lines. Each path from the root of the binary three (at the top) to any of its nodes, corresponds to a segment (with the same colour) in the sunburst (at the bottom) and it represents a tumour subpopulation with an associated genomic signature (also reported in table B). The branches of the tree reflect the recursive steps of the algorithm with colored arrows specifying a recursion that aims at refining the analysis of the subpopulation corresponding to the source node (green) or at analysing its complementary subpopulation (orange). The prevalence of the identified signatures, and their hierarchical co-occurrence is represented by the length of the corresponding segment in the sunburst. **B.** Cell Line Map table including microsatellite instable cell lines mirroring the genomic signatures defining tumour subpopulations of the CELLector search space. The models in green represent a possible choice of 4 cell lines that could be selected in the presented case study.

To facilitate mapping cell lines onto patient segments and their prioritisation, the search space building algorithm also stores intermediate and final results in a binary tree structure. Particularly, each genomic alteration (or alteration pair/triplet) identified recursively through the search space building algorithm is stored in a tree node. Linking nodes identified in adjacent recursions yields a binary tree, which provides an alternative way of visualising *the CELLector search space*, better reflecting its construction steps (Figure 2A).

This tree defines, by construction, logic formulas, one for each patient segment. In fact, each individual path (from the root to any node) of the tree corresponds to a segment in the sunburst visual (Figure 2A), and it defines a rule (a logic formula, or signature). This is represented as a logic AND of multiple terms, one per each node in the path (Figure 2B). These terms are negated (∼) when the corresponding node is linked through the path to another node mapping alteration(s) identified in a complement recursion step rather than a refinement recursion step.

Per construction, if the genome of a given patient of the analysed cohort satisfies the rule associated to a given path in the tree, then that patient belongs to the segment associated with that path, or (for simplicity) to its terminal node. Similarly, cell lines are mapped onto patient segments. Collectively, all the paths in the search space provide a representation of the spectrum of molecular alterations (and their combinations) most frequently observed in a given cancer type, and their clinical prevalence in the analysed cohort of patients (Figure 2A).

The order of the nodes resulting from a visit of this tree detailed below, defines the priority of the corresponding mapped cell lines. This provides a possible choice of the best *n* cell lines, maximising the covered genomic heterogeneity of the considered cohort of cancer patients.

Once constructed, the CELLector search space is mined by a second module selecting the most representative set of cell lines maximising their covered genomic heterogeneity, via a guided visit of the search space tree (STAR Methods), thus also identifying tumour subtypes lacking representative cell line models (Figure 2B). Particularly, this visit starts from the centre of the search space sunburst considering the largest and innermost segment (corresponding to the genomic alteration - or set of alterations - most frequently occurring in the analysed cohort of patients), and proceeds through adjacent segments (from the 2nd largest one to the 3rd and so on). When all the segments in a given level have been visited and stored in a queue, the algorithm removes the first segment from the queue and the visit restarts considering its sub-segments in the outer level of the sunburst, from the largest one to the 2nd largest one and so on. The recursion continues until the queue is emptied. Each time a segment is visited for the first time, the algorithm picks one of the cell lines mapped on it. The resulting ordered list of cell lines is outputted as a possible optimal selection as it includes cell lines that are representative of the largest subset of patients (Figure 2B).

#### CELLector capabilities

CELLector can assist the selection of the best-representative preclinical models to be employed in molecular oncology studies. It also enables a frequency-based molecular subtyping/classification of any disease cohort. As detailed in the previous section, one of the approaches that users can pursue with CELLector is a simple *guided visit* of its search space to select the optimal set of *n* cell lines to be included in a small-scale *in vitro* study or a low-throughput screen. The selected cell lines are picked from those mapped to the first *n* nodes of the searching space, as they appear in the guided visit of the corresponding tree/sunburst and, per construction, this guarantees that the coverage of the genomic heterogeneity of a particular cancer type is maximised by the selected cell lines.

Another functionality of CELLector is the provision of a quantitative estimation of the *quality* of a given cell line in terms of its ability to represent the entire cohort of considered patients. This is quantified as a trade-off between two factors. The first factor is the length of the CELLector signatures (in terms of number of composing individual alterations) that are present in the cell line under consideration. This is proportional to the granularity of the representative ability of the cell line, i.e. the longest the signature the more precisely defined is the represented sub-cohort of patients. The second factor is the size of the patient subpopulation represented by the signatures that can be observed in the cell line under consideration, thus accounting for the prevalence of the sub-cohort modeled by that cell line.

In addition, given that the choice of appropriate *in vitro* models often depends on the context of the study, while constructing the search space users can restrict the analysis to a given sub-cohort of patients, by determining a priori based on the presence/absence of a given genomic feature. For example, users can restrict the CELLector analysis to subset of tumours harbouring *TP53* mutations, or genomic alterations in the PI3K/Akt signalling pathway. In this case, only tumours characterised by these features are taken into consideration when building the CELLector search space (as detailed in the next section and in Supplemental Information - Case Studies). Notably, this can also involve other user-defined characteristics, for example the microsatellite instability (MSI) status of cancer cell lines. As a consequence, CELLector allows users to flexibly tailor the selection of cell lines in a context-dependent manner.

The CELLector R package and R Shiny app are both interfaced with the Cell Model Passports database of cancer models (van der Meer et al., 2019) and can reassemble the built-in genomics datasets (in a fully customisable way) to synchronise them to their most recent releases. In particular, this is accomplished by a separate module: the Binary Event Matrix (BEM) builder, which can be also used to assemble and export genomic binary matrices (for both cell lines and primary tumours). These can be used within CELLector itself or by other tools, for instance to identify markers of drug responses or gene essentiality using publicly available data for example, from the Genomics of Drug Sensitivity in Cancer (GDSC) portal (Iorio et al., 2016; Yang et al., 2012; Garnett et al., 2012) or the Cancer Dependency Map web-sites (Behan et al., 2019; Tsherniak et al., 2017), and tools such as GDSCtools (Cokelaer et al., 2018), among others.

Finally, the CELLector R Shiny app provides additional functionalities enabling an interactive exploration of the tumour/cell line genomic features and final results. A tutorial demonstrating all these functionalities, example case studies, and a step-by-step guide to reproduce the results reported in these case studies is provided as Supplemental Information.

### Use Case: Selecting Microsatellite Instable Cell Lines Representing *BRAF* Mutant Colorectal Cancers

In this section, we present a practical example to demonstrate the usefulness of CELLector in an experimental study design. Detailed instructions on this and other use cases are provided in the user tutorial available as Supplemental Information. In this example, we want to identify the most clinically relevant microsatellite instable cell lines that capture the genomic diversity of a sub-cohort of colorectal cancer patients that harbour *BRAF* mutations. *BRAF* mutant colorectal cancers have a low prevalence (5%-8%) and very poor prognosis (Sanz-Garcia et al., 2017). In this example, the model selection will be performed accounting for somatic mutations that are prevalent in at least 5% of the considered colorectal patient cohort (Figure 2A: box 1 and box 2).

#### Building the CELLector search space

After setting the CELLector app parameters to reflect the search criteria detailed in the previous section (Figure 2A: box 1 and box 2), the CELLector search space is assembled using a built-in dataset containing the genomic characterisation of a cohort of 517 colorectal cancer tumours (Supplemental Information Table S1 and STAR Methods).

First, the cohort is reduced to the tumours harbouring *BRAF* mutations (*n=86*, Figure 2A: node 1). CELLector then identifies 3 major molecular subpopulations characterised, respectively, by *APC* mutations (Figure 2A: node 2), *FBXW7* mutations (Figure 2A: node 3), and *PIK3CA* mutations (Figure 2A: node 10), collectively representing 85% of the *BRAF* mutant cohort. The remaining 15% of *BRAF* mutant tumours do not fall into any of the identified molecular subpopulations, i.e. they do not harbour *APC, FBXW7* nor *PIK3CA* mutations; Figure 2A).

The largest molecular subpopulation (58.14%, harbouring *BRAF* and *APC* mutations) is assigned to the root of the search space (Figure 2A: node 2, in purple). The second largest subpopulation (16.28%) is characterised by the co-occurrence of *BRAF* and *FBXW7* mutations in the absence of *APC* mutations (Figure 2A: node 3, in magenta), and the third largest subpopulation (10.47%) harbours the *BRAF* and *PIK3CA* mutations in the absence of both *APC* and *FBXW7* mutations (Figure 2A: node 10, in cyan). At this point, each identified tumour subpopulation is further refined based on the prevalence of remaining set of alterations (STAR Methods). This process runs recursively and stops when all alteration sets with a user-determined prevalence (in this case 5%, Figure 2A: box 1) are identified. In this study case, a total number of 10 distinct tumour subpopulations with corresponding genomic signatures are identified (Figure 2).

#### Selection of representative in vitro models

The CELLector search space generated as detailed in the previous section is next translated into a Cell Line Map table (Figure 2B), indicating the order in which cancer *in vitro* models mirroring the identified genomic signatures should be selected, and accounting also for tumour subpopulations currently lacking representative *in vitro* models. This selection order is defined by a guided visit of the CELLector search space (introduced in the previous sections and detailed in the STAR Methods), aiming at maximising the heterogeneity observed in the studied primary tumours. The Cell Map table uncovers the complete set of molecular alterations (e.g. genomic signatures) defining each tumour subpopulation (in this example *n=10*). For example, a *BRAF* mutant colorectal tumour subpopulation (node 8, 9.30% of tumours) is characterised by the co-occurrence of *BRAF*, *APC, PIK3CA, PTEN, TP53 and KRAS* mutations; this genomic signature (*BRAFmut APCmut PIK3CAmut PTENmut TP53mut KRASmut)* is not mirrored by any of the available microsatellite instable colorectal cancer models included in the built-in dataset. On the contrary, the subpopulation characterised by the co-occurence of BRAF, APC and TP53 mutations in absence of PIK3CA mutations (*BRAFmut APCmut ∼PIK3CAmut TP53mut*, node 5, 15.12% of tumours) is represented by microsatellite instable KM12 and LS-411N cell lines (Figure 2B).

Finally, representative cell lines are picked from each of the molecular tumour subpopulations (as detailed in the STAR Methods) starting from the most prevalent one, i.e. as they appear in the Cell Line Map table. A possible choice of *in vitro* models that best represent the genomic diversity of the studied tumour cohort include: LS-411N, SNU-C5, RKO and KM12 (Figure 2B shown in green). Additional case studies are included in the user tutorial provided as Supplemental Information.

### CELLector Bridges Cancer Patient Genomics with Cell Line Based Pharmacogenomic Studies

To fully demonstrate the potential of the CELLector analytical framework we performed two landmark analyses: (i) linking results and findings from large scale drug screens performed *in vitro* to cancer patient cohorts; (ii) performing a systematic estimation of the largest patient sub-cohorts currently lacking representative *in vitro* models across multiple cancer types (Figure 3). Executing these analyses routinely and on increasingly larger tumour datasets in the future might serve to (i) *in-silico* prescribe drugs to precisely defined sub-cohorts (segments) of cancer patients; (ii) prioritise *in vitro* models for future development.

**Figure 3.**
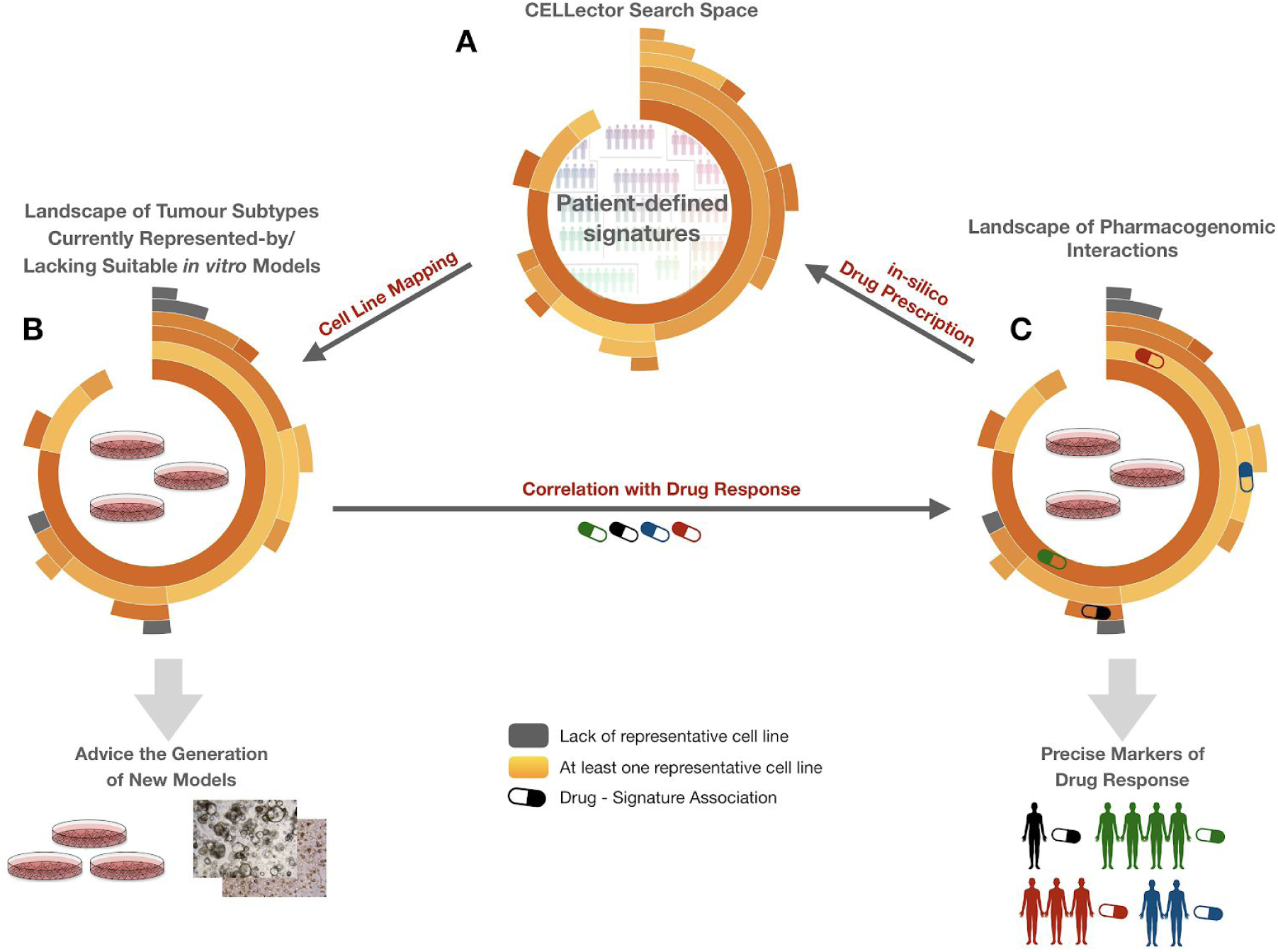
CELLector bridges cancer patient genomics with cell line based pharmacogenomic studies. **A.** CELLector provides patient-defined signatures, which are mapped to tissue-matched cell lines. **B.** This enables to assemble a landscape of tumour subtypes currently represented-by (orange)/lacking (grey) suitable *in vitro* models, which represents a valuable resource to prioritize the generation of new *in vitro* models across different cancer types. **C.** The patient-defined signatures can then be used as possible predictors of differential drug sensitivity in cancer cell lines providing an estimation of potential clinical relevance and size of the patient cohort that would respond to the therapy. This framework provides a means for *in silico* prescribing drugs to precisely defined subpopulation of cancer patients.

Particularly, by making use of publicly available data (Iorio et al., 2016), we applied CELLector systematically to a large number of patient cohorts (STAR Methods) which were segmented into patient subtypes with associated combinations of the most prevalent genomic alterations, e.g patient-defined signatures across different cancer types (Figure 3A). Then we mapped cell lines onto the identified patient segments based on the collective presence/absence of the corresponding signatures. In this way we generated a landscape of cancer patient sub-cohorts currently represented/non-represented by available *in vitro* models, which can be used as a rule-book for the generation of new cell lines and organoid models (Figure 3B), as well as a means to directly link cancer patient subpopulations to large cell line based pharmacogenomic studies (Figure 3C). In fact, the identified CELLector signatures that are present in a suitable number of cancer cell lines can be systematically correlated with drug responses observed in large scale cell line based screenings, thus providing a powerful and clinically relevant way to associate tumour genotypes (together with their prevalence) with established or potential cancer therapies. As we show in the following sections, this enables an *in silico* prescription of existing drugs directly to precisely defined subgroups of cancer patients, and might serve for the identification of complex and more robust markers of drug response (Figure 3C).

#### Landscape of tumour subtypes currently represented-by/lacking suitable in vitro models

To estimate the genomic heterogeneity of primary tumours and to assemble a landscape of patient subtypes currently represented-by/lacking suitable *in vitro* models, we systematically applied CELLector to large cohorts of primary tumours across 22 cancer types, focusing on cancer driver somatic mutations (SMs), copy number alterations (CNAs) and combination of both SMs and CNAs, occurring in at least 2% of a patient cohort (Figure 4A, Supplementary Figure S1, Supplementary Data S1). This analysis identified a total number of 718 patient-defined signatures (and corresponding patient segments) and highlighted that 46.8% (n=336) of them are covered by at least one tissue-matched cell line in the Cell Model Passport collection (a widely representative collection of *in vitro* models) (van der Meer et al., 2019). Strikingly, the remaining 53.2% (n=382) of identified patient segments (signatures) lack representative *in vitro* models (Figure 4B, Supplementary Figure S2, Supplementary Data S1).

**Figure 4.**
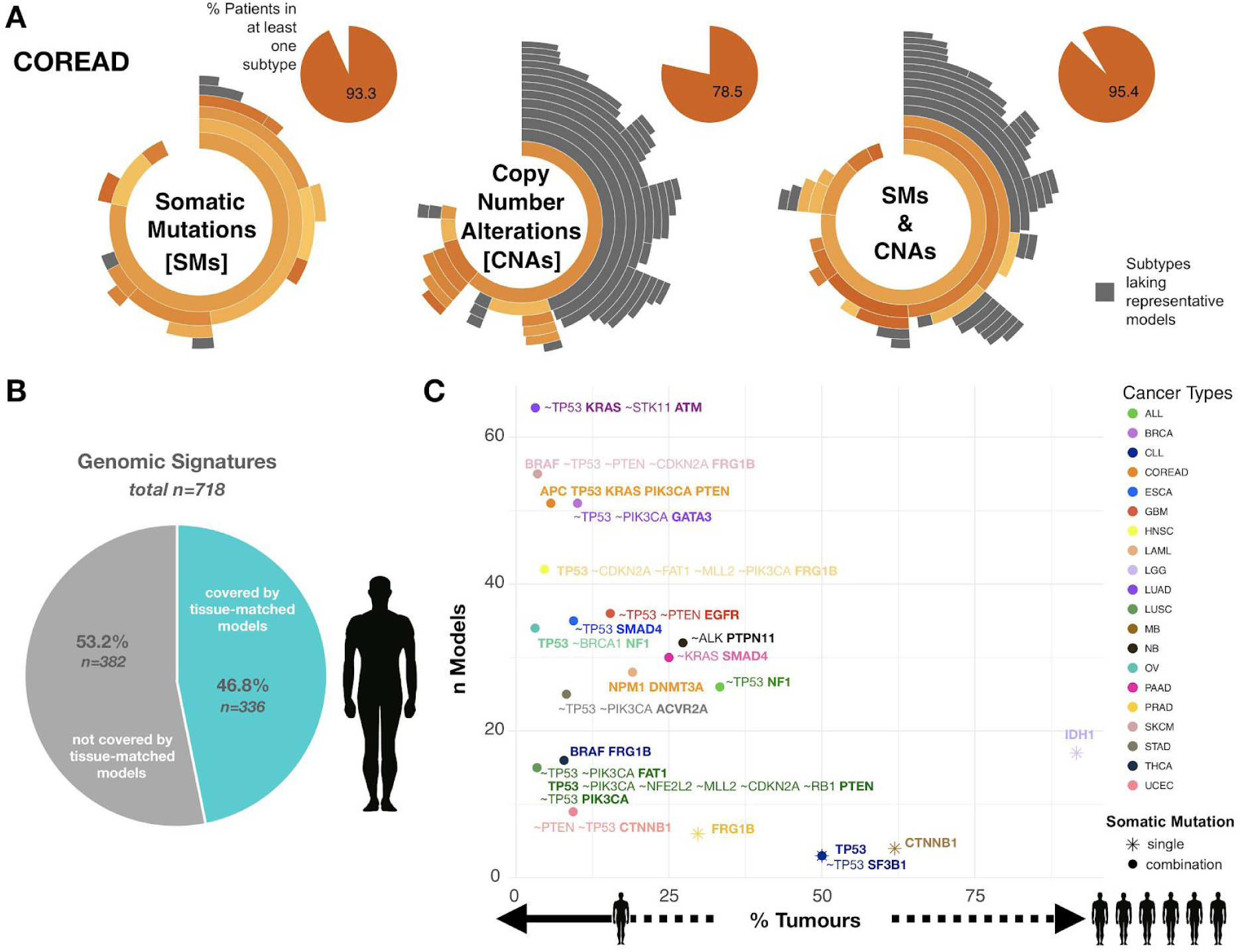
Systematic identification of patient populations lacking representative *in vitro* models. **A.** Landscape of tumour subtypes represented-by/lacking suitable *in vitro* model for an example of cancer type (colorectal adenocarcinoma - COREAD) across different genomic data types. Grey sunburst segments represent tumour subpopulations not captured by any of the tissue-matched cell lines, while orange segments represent COREAD subpopulations represented by at least one tissue-matched model. Results for all the other analysed cancer types are shown in Supplementary Figure S1. **B.** Patient-defined CELLector signatures prevalent in at least 2 % of patient cohort across 22 cancer types. Pie chart represents a summary of the systematic CELLector analysis across different genomic data types (somatic mutations, copy number alterations and combinations of both somatic mutations and copy number alterations). **C.** Missing models in demand: each point represents the most prevalent patient subpopulation that is not captured by tissue-matched cell line when somatic mutations prevalent in at least 2% of patient cohort are considered; x-axis indicates percent of considered patient cohort, y-axis indicates the total number of considered tissue-matched cell line models in respective analyses.

Collectively, the tumour subtypes defined by the signatures and covered by at least one tissue-matched model spanned across 56% of patients with available data when considering either signatures of mutations or copy number alterations (74% when considering both); 35% and 32% (respectively for mutations, and copy number alterations, 14% when considering both) of patients were not included in any of the CELLector defined sub-cohorts, indicating that their genomes only host rare mutations/CNAs (frequency < 2% of their cohort). Finally, 8% and 12% (respectively for mutations, and copy number alterations, 11% when considering both) of patients fall into at least one recurrent subtype (covering > 2% of patients) strikingly lacking representative *in vitro* models (Supplementary Figure S3A). Among the most underrepresented cancer types we found brain lower grade glioma (LGG; 95.1% of patients falling into at least one subgroup lacking representative *in vitro* models), followed by prostate adenocarcinoma (PRAD; 62.16% of patients) (Supplementary Figure S1).

This analysis also highlighted large disease subtypes that should be taken in consideration while prioritizing new *in vitro* models for future development (Figure 4C). As an example, despite the large availability of LGG cell lines (when compared to other cancer types), none of these harbour *IDH1* mutations, which define the largest LGG patients subpopulation (91.6% tumours). Similarly, there are no medulloblastoma (MB) cell lines harbouring *CTNNB1* mutations, defining 61.9% of MB patients. Lung adenocarcinoma (LUAD) is one of the cancer types with the largest number of available cell lines, however 3.17% LUAD patients that harbour mutations in *KRAS* and *ATM* in the absence of *TP53* and *STK1* mutations (*∼TP53mut, KRASmut, ∼STK11mut, ATMmut*) are not represented by any tissue-matched *in vitro* model (Figure 4C).

#### CELLector identifies clinically relevant tumour subtypes and enables precise in silico drug prescriptions

Literature mining confirms both prevalence and clinical relevance of some of the genomic signatures identified by CELLector. For example, some of the CELLector signatures identified in the *BRAF*-mutant colorectal tumours case study (shown in the previous section) are linked to differential prognosis in colorectal tumour stratification (Schell et al., 2016). Specifically, co-occurrence of mutations in *APC*, *TP53* and *KRAS* (defining the *triple mutant AKP subtype*), and the co-occurrence of *APC* and either *KRAS* (*AK subtype*) or *TP53* (*AP subtype*) mutations are identified by CELLector when performing a COREAD specific analysis using mutation data with default parameters (Supplementary Figure S3B, details in Case Study 2 of Supplemental Information). All these subtypes are reported to have a discriminative prognostic power in colorectal tumour stratification (Schell et al., 2016). As another example, the genomic landscape of mutually exclusive *BRAF*- and *NRAS*-mutant melanomas detected by CELLector when performing a SKCM specific analysis with default parameters can be further contextualized, providing insights into co-occurring mutations and copy number alterations (Hodis et al., 2012).

To programmatically estimate the potential clinical relevance and translational potential of the CELLector signatures, we systematically correlated their status observed in the cell lines with their responses to 495 compounds using publicly available data from the GDSC project (Garnett et al., 2012; Iorio et al., 2016, Picco et al., 2019). Furthermore, we also performed a systematic survival analysis using CELLector signatures and individual CFEs to define subpopulations of patients to be contrasted for differential prognosis using data from cBioPortal (Cerami et al., 2012).

##### CELLector signatures are involved in novel robust pharmacogenomics interactions

We performed cancer-type specific Analyses of Variance (ANOVAs) to test if patient-defined combinations of genomic functional events defined by the CELLector signatures correlate with the responses to any of 495 drugs screened in cancer cell lines (using data from Picco et al., 2019), and whether they yield more robust pharmacogenomic associations when compared to those involving individual genomic features, or cancer functional events, CFEs, as defined in Iorio et al., 2016.

To this aim, we computed CELLector signatures for 13 cancer types, considering somatic mutations, copy number alterations and methylation of gene promoters occurring in at least 2% of the patient cohort under consideration (Supplementary Data S2), and used these as factors together with individual CFEs (Supplementary Figure S4A, Supplementary Data S3) in systematic cancer-type specific ANOVAs (similarly to the analyses performed in (Garnett et al., 2012; Iorio et al., 2016)). We considered only CFEs and CELLector signatures with at least 3 positive/negative available cell lines for a total number of 498 unique CFEs (median across cancer types = 50, involved in a total number of 264,479 tests across cancer types and drugs) and 224 unique CELLector signatures (median = 13, involved in 88,523 tests, Supplementary Figure S4A).

Results encompassed 349 significant differential drug responses (*p < 0.001, FDR < 25, Cohen’s d > 1*) spanning across 12 cancer types (Figure 5A), and involving 197 unique drugs and 148 unique tested features (Supplementary Figure S4B). Of these, 93 associations resulted from tests where the underlying factor was a CELLector signature (median = 7 per cancer type) and 256 from tests involving individual CFEs (median = 11 per cancer type, Figure 5A, Supplementary Figure S4B, Supplementary Data S4). Importantly, 38 drug-signature associations involved 25 unique signatures and 34 unique drugs not significantly associated with any individual CFEs (Figure 5A, Supplementary Figure S4B) thus demonstrating the ability of the CELLector signatures to uncover new pharmacogenomic interactions.

**Figure 5.**
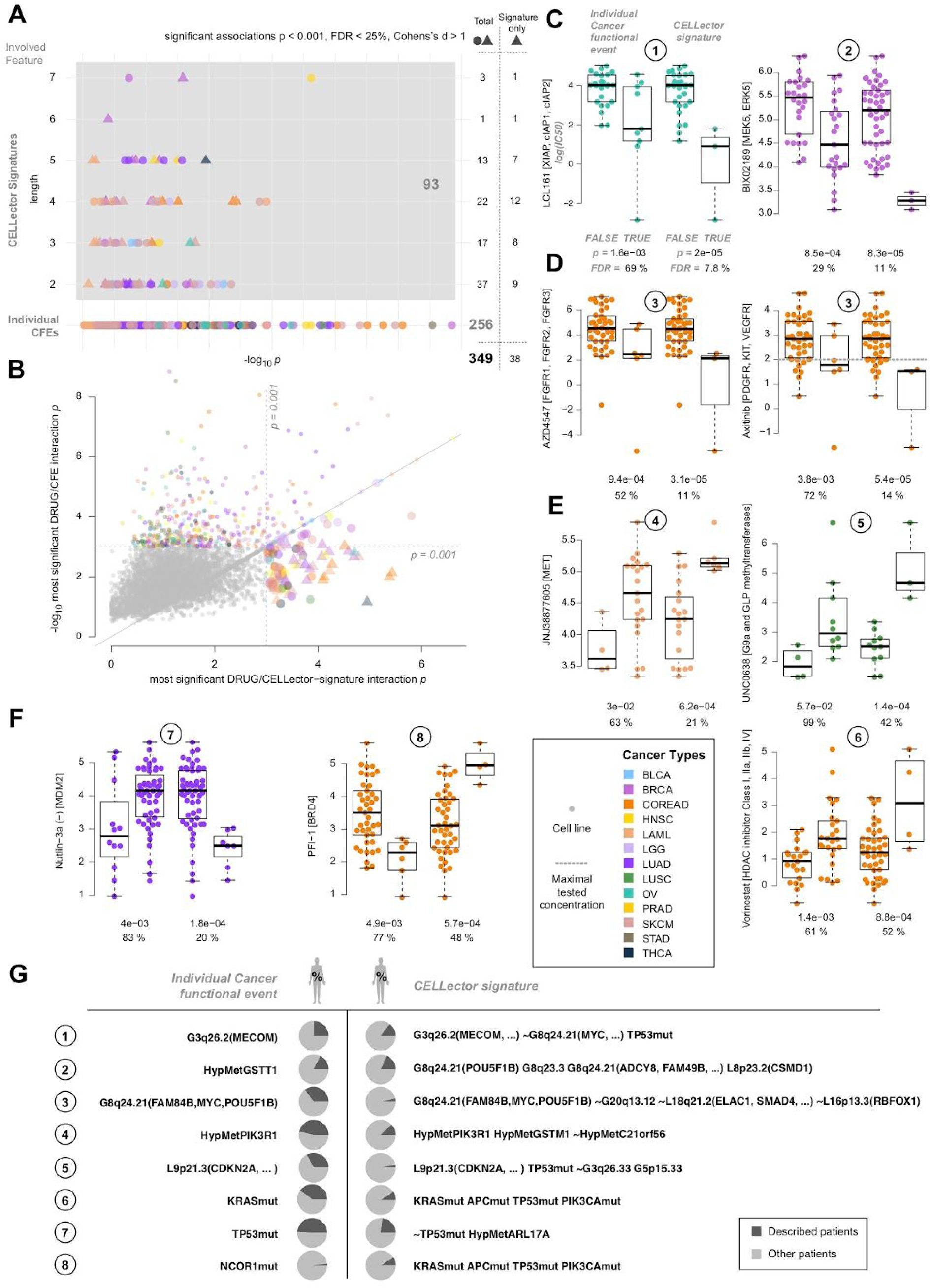
Patient-defined CELLector signatures improve pharmacogenomic studies and uncover new pharmacogenomic interactions. **A.** Overview of significant pharmacogenomic interactions (data points) across cancer types and types of involved features: respectively, individual cancer functional events (CFEs, on the first row) and CELLector signatures (with coordinates on the y-axis indicating the signature length, in terms of contained CFEs). Coordinate on the x-axis indicate the interaction significance. Point colors indicate cancer type specific analysis in which an interaction was detected as significant and triangular shapes indicate drug-signature interaction involving a drug with no significant drug-CFE interactions. **B.** P-value comparison for top significant drug-signature associations and drug-CFE associations for all screened drugs and across cancer types, as indicated by the different color. Each point represents a drug with coordinates on the two axis reflecting the significance level of the top significant associations. **C.** Pairs of plots with examples of drug-signature associations (second plot in each pair) that are much more significant than the top significant drug-CFE association involving the same drug (first plot in each pair). Each circle represents a cell line with coordinate on the y-axis indicating the log IC50 of the drug specified in the y-axis label. In each individual plot, cell lines are partitioned into two groups based on the status of a genomic feature (TRUE or FALSE, indicating respectively the presence or absence of that feature). These features can be individual CFEs (first plot in each pair) or CELLector signatures (second plot in each pair), and are specified in the table in G. P-values and False Discovery Rates are from Analysis of Variance (ANOVA) tests assessing the extent of differential drug response across the considered dichotomies of cell lines. The boxes cover interquartile ranges with median lines drawn within them. Whiskers extend to a maximum of 1.5 times the size of the interquartile range. Point colors indicate the cancer type of origin of the cell lines. **D.** As for C but showing examples where CELLector signatures including MYC amplifications together with other CFEs (or their negation) are associated with drug response more significantly than MYC amplifications alone. **E.** As for C but showing pharmacogenomic associations involving increased drug resistance rather than sensitivity. **F.** As for C but showing examples where a CELLector signatures define subgroups of very sensitive (respectively resistant) cell lines to a drug with an associated CFE marker of resistance (respectively sensitivity). **G.** Features (CFEs and CELLector signatures) whose status is used in the pair of plots in CDEF to dichotomised the cell lines and contrast their IC50s. The numerical labels on the left realise the mapping between each row and each pair of plots. Pie charts indicate percentage of patients harboring the indicated CFE (left panel) or CELLector signature (right panel), and whose cancer type matches that indicated by the color in the plot pairs.

Furthermore across all cancer types, we observed that for 88 unique drugs at least one significant drug-signature association (*p* < 0.001) was more significant than the most significant drug-CFE association involving the same drug (Figure 5B, Supplementary Figure S5A, Supplementary Data S5), in at least one cancer-type specific ANOVA. This was observed invariantly across cancer types (with the exception of the PRAD specific analysis) with a median number of 4 drugs per cancer type (Figure 5B, Supplementary Figure S5A).

For several cases the difference in significance was conspicuously large and associated with an increased level of drug sensitivity (Figure 5CD and Supplementary Figure S5B). For example, we found that increased sensitivity to the second mitochondrial-derived activator of caspases (*SMAC*) mimetic and inhibitor of *IAP* (Inhibitor of Apoptosis Protein) LCL161 is weakly associated (*p = 1.6 x 10*^-3^*, FDR = 69%*) with gains of a genomic segment containing the *MDS1* and *EVI1* complex locus (MECOM) oncogene in OV cell lines (occurring in 25% of patients). A signature occurring in 15% of cancer patients and accounting for amplifications of the same segment, in combination with TP53 mutations and copy-number wild-type MYC is associated with sensitivity to LCL161 much more significantly (*p = 2 x 10*^-5^*, FDR = 7.8%*, Figure 5C). Further, gains of three genomic segments of chromosome 8, containing the POU Class 5 Homeobox 1B (POU5F1B), the Adenylyl cyclase type 8 (ADCY8), and other genes, in combination with losses of CUB and Sushi Multiple Domains 1 (CSMD1) - all occurring simultaneously in 18% of BRCA patients - was much more significantly associated with sensitivity to the *MEK5/ERK5* inhibitor BIX02189 (*p = 8.3 x 10*^-5^*, FDR = 11%*, Figure 5C) than the top predictive individual CFE (hypermethylation of the glutathione S-transferase theta 1 - GSTT1 - gene promoter, *p = 8.5 x 10*^-4^*, FDR = 29%*, occurring in 17% of BRCA patients).

Notably, in many other cases, CFEs consisting of mutations or amplifications of canonical oncogenes associated with increased sensitivity to a given drug (often targeting their coded protein) with a lower significance than a drug-signature association involving the same CFEs in combination with secondary genomic alterations (Figure 5D and Supplementary Figure S5B). For example, in COREAD cell lines, MYC amplifications (occurring in 35% of COREAD patients) were weakly associated with increased sensitivity to the *FGFR* inhibitor AZD4547, the *PDGFR/KIT/VEGFR* ihibitor Axitinib, the *VEGFR* inhibitor Tivozanib, and the inhibitor of *TERT* BIBR-1532 (*p < 0.05*, FDRs ranging in [52%, 93%], Figure 5D and Supplementary Figure S5B). Nevertheless, increased sensitivity to these compounds was much more significantly associated with a signature composed of MYC amplifications in combination with CN wild-type segments containing, among other genes, the RNA Binding Fox-1 Homolog 1 (RBFOX1), the *SMAD* family member 4 (SMAD4) and the ElaC Ribonuclease Z 1 (ELAC1) - (max *p = 5 x 10^-4^*, FDR ranging in [11%, 45%], Figure 5D and Supplementary Figure S5B): a signature observed in 3% of COREAD patients. Among the other examples worthy of note (reported in Supplementary Figure S5B), in SKCM cell lines a signature accounting for BRAF mutations and hypermethylation of the Zinc Finger Protein 714 gene promoter (observed simultaneously in 26% SKCM patients) was more significantly associated (*p = 1.3 x 10^-4^, FDR = 15%*) with sensitivity to *LIM* Domain Kinase 1 (LIMK1) inhibitor BMS4 than BRAF mutations considered alone (*p = 5.7 x 10^-4^, FDR = 23%*, Supplementary Figure S5B), observed in 51% of patients.

We observed similar outcomes when looking at drug resistance associations, with several drugs showing a CELLector signature as top significant hit. For example in AML cell lines, the combined promoter hypermethylation of the Phosphoinositide-3-Kinase (*PI3K*) Regulatory Subunit 1 (PIK3R1) and the glutathione S-transferase (GSTM1) genes in the absence of the hypermethylation of the Spermatogenesis And Centriole Associated 1 Like (C21orf56) genes - a signature observed in 12% of patients - was robustly associated with resistance to the MET inhibitor (*p = 6.2 x 10^-4^, FDR = 21%*, Figure 5E). As for the previous examples, this association was much more significant than the top significant one at the level of individual CFEs, i.e. hypermethylation of the PIK3R1 promoter alone (*p = 3 x 10^-2^, FDR = 63%*) - observed in 47% patients - downstreaming of c-Met and with an established role in the autocrine activation of AML cells (Kentsis et al., 2012).

As for the sensitivity markers, CFEs consisting of altered canonical oncogenes/tumour-suppressors that were associated with increased resistance to a given drug had their predictive ability improved when they were considered together with secondary genomic alterations in a CELLector signature (Figure 5E). For example in LUSC cell lines, losses of the cyclin-dependent kinase Inhibitor 2A (CDKN2A) - observed in 33% of patients - were weakly associated with increased resistance to the G9a and GLP methyltransferases inhibitor UNC0638 (*p = 0.057, FDR = 99%*). Differential resistance to this drug raised significantly (*p = 1.4 x 10^-4^, FDR = 42%*, Figure 5E) when contrasting the responses of cell lines based collectively on the presence of CKDN2A losses, TP53 mutations, gain of another segment on chromosome 5 not containing any known cancer genes and the CN wild-type status of a segment on chromosome 3 (a signature observed in 3% of patients). Furthermore, in COREAD cell lines, a signature made of mutations in APC, TP53, KRAS and PIK3CA (observed in 9% of COREAD patients) was more robustly predictive of resistance to the multiclass histone deacetylase inhibitor Vorinostat (*p = 8.8 x 10^-4^, FDR = 52%*), than KRAS mutations alone (*p = 1.4 x 10^-3^, FDR = 61%*) - observed in 40% of patients (Figure 5E).

Furthemore, our analyses unveiled potential novel multivariate markers able to define very precisely sub-cohorts of putative sensitive (resp. resistant) patients in the context of drugs with established resistance (resp. sensitivity) markers (Figure 5F). For example, in LUAD cell lines, our analyses confirmed (although weakly) the established association between TP53 mutations (observed in 49% of LUAD patients) and resistance to the inhibitor of the *TP53-MDM2* interaction Nutlin-3a (Kojima et al., 2006) (*p = 1.4 x 10^-3^, FDR = 61%*). At the same time, a much more significant associations (*p = 1.8 x 10^-4^, FDR = 20%*) between sensitivity to Nutlin-3a and a CELLector signatures describing a sub-cohort of LUAD patients (23%) with wild-type TP53 and hypermethylation of the ADP Ribosylation Factor Like GTPase 17A (ARL17A) gene promoter was unveiled (Figure 5F). Viceversa, a signature describing a sub-cohort of COREAD patients with mutations in APC, TP53, KRAS, and PIK3CA (9%) was associated with resistance to the inhibitor of the bromodomain-containing protein 4 (BRD4) PFI−1 more significantly (*p = 5.7 x 10^-4^, FDR = 48%*) than mutations in the Nuclear Receptor Corepressor 1 (NCOR1) gene, observed in 2% of patients and the most predictive individual CFE for this drug in COREAD cell lines, associated with increased sensitivity (*p = 4.9 x 10^-3^, FDR = 77%*, (Figure 5F).

Finally, CELLector signatures often improved the significance of a CFE/drug association providing the basis for mechanistic interpretations and potentially refining patient stratification for precision medicine. For example, our analyses detected weakly significant associations between *KRAS*-mutations and increased sensitivity to the *MEK1/2* inhibitor Selumetinib (p = 0.04, FDR = 97%, Figure 6A) and Dabrafanib, a BRAF inhibitor (p = 0.001, FDR = 54%, Figure 6A), in LUAD cell lines. We found that two complementary CELLector signatures were much more significantly associated respectively with sensitivity to Selumetinib (p = 3.3 x 10^-4^, FDR = 29%) and Dabrafenib (p = 8.4 x 10^-4^, FDR = 51%). These signatures substratify *KRAS*-mutant LUAD patients (25% of the LUAD cohort considered in this study), based (the first) on the presence of TP53 mutations and ARL17A promoter hypermethylation (observed with KRAS mutations in 7% of the cohort) and (the second) based on the absence of these two alterations and hypermethylation of the GSTT1 promoter (observed in 5% of the cohort) (Figure 6A).

**Figure 6.**
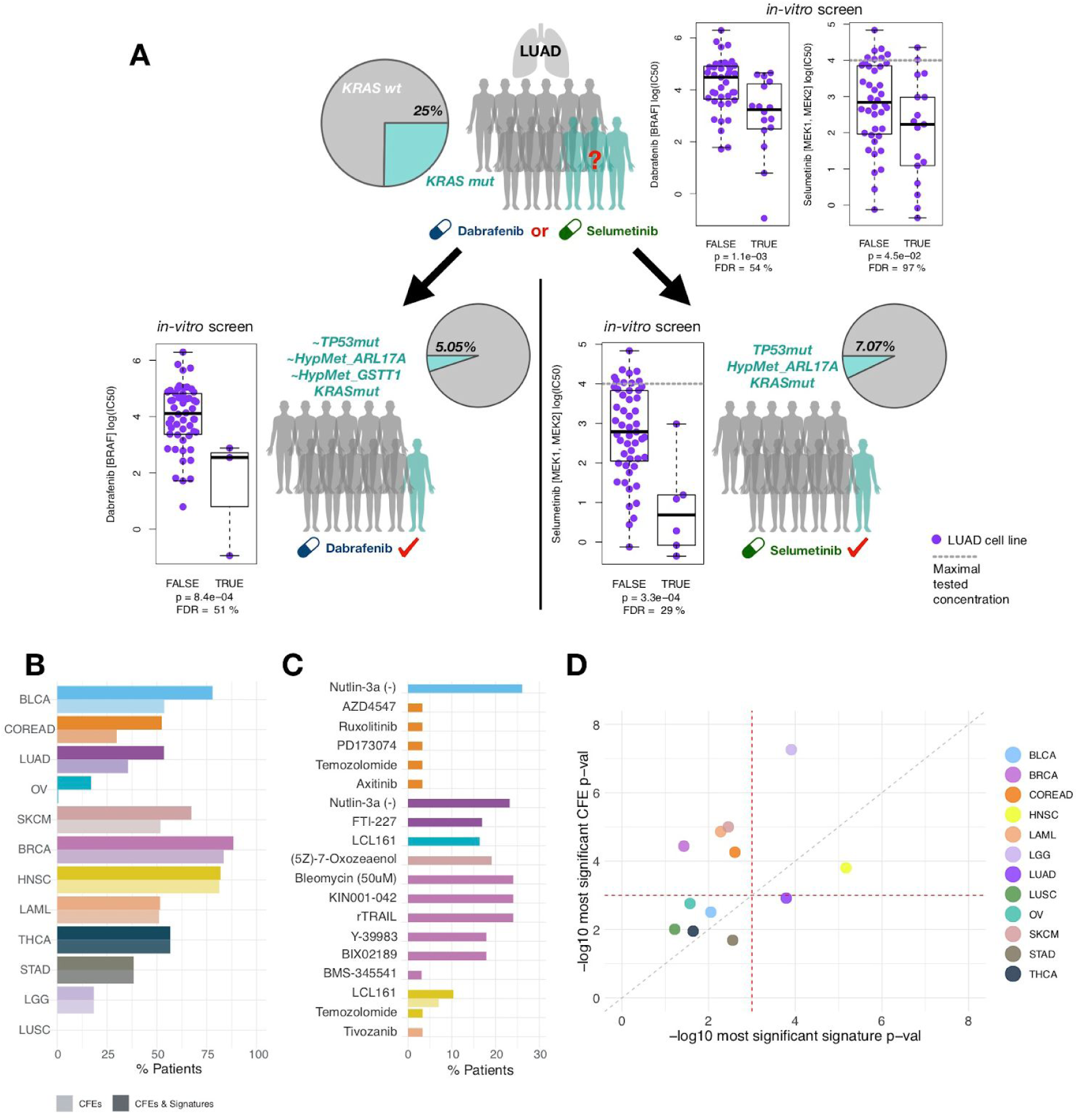
CELLector signatures enhanced precision medicine landscape. **A.** In vitro pharmacogenomic data weakly confirms an association between KRAS mutations and sensitivity to inhibitors of the *MAPK* pathway inhibitors (Dabrafenib and Selumetinib) in Lung Adenocarcinoma (LUAD). Two patient-derived CELLector signatures account for the mutational status of KRAS with added the presence/absence of supplemental TP53 mutations and hypermethylation of the ARL17A gene promoter. These signatures describe two complementary sub-cohorts of the KRAS mutant LUAD patients and cell lines mapped by CELLectors onto these two sub-cohorts are much more significantly sensitive Dabrafenib and Selumetinib, respectively. Thus the two drugs could be prescribed *in silico* to these two distinct patient sub-cohorts. Pie-charts show KRAS mutation and CELLector signature prevalence in LUAD patients. In the plots, each circle represents a cell line with coordinate on the y-axis indicating the log IC50 of the drug specified in the y-axis label. Cell lines are partitioned into two groups based on the status of KRAS (TRUE or FALSE, indicating respectively the presence or absence of mutations) or the CELLector signatures (TRUE or FALSE depending on the logic formula described in the signature being satisfied or not). P-values and False Discovery Rates are from Analysis of Variance (ANOVA) tests assessing the extent of differential drug response across the considered dichotomies of cell lines. The boxes cover interquartile ranges with median lines drawn within them. Whiskers extend to a maximum of 1.5 times the size of the interquartile range. **B.** Percentages of patients harboring a sensitivity marker, either an individual Cancer Functional Event (CFE, brighter colour) or both CFEs and CELLector signatures (darker colour) for at least one of the screened drugs considered in our ANOVAs, across cancer types (as indicated by the different colours). **C.** Examples of drugs for which the percentages of patients harboring a sensitivity marker increases when considering signatures in addition to CFEs, across cancer types. **D.** Systematic comparison of differential survival Cox p-values (corrected by age and gender) for top predictive individual CFE and CELLector signatures across cancer types.

Finally, to evaluate the potential clinical impact of the CELLector signatures, we quantified, across cancer types, to what extent considering drug-signature sensitivity associations in addition to drug-CFE sensitivity associations varies the number of patients whose cancer hosts at least one drug sensitivity marker, compared to considering CFE-drug associations only. To this aim we quantified the percentages of patients, across cancer types, harboring at least one feature significantly associated with increased drug response (p < 0.001, FDR < 25%, for at least one drug), with a feature being a CFE in the first case and a CFE or a CELLector signature in the second case.

Notably, the number of patients harbouring sensitivity markers increased when considering both CFEs and CELLector signatures, for 8 out of 12 cancer types (Figure 6B, median = 4.88%, ranging from 24% for BLCA to 0% for THCA, STAD and LGG).

Specific sets of drugs explain these differences (Figure 6C), including 18 unique drugs with novel CELLector signature markers of drug sensitivity. These encompassed targeted compounds (such as Nutlin-3a in BLCA and LUAD, RTK signaling pathways inhibitors AZD4547, PD173074, and Axitinibin in COREAD and Tivozanib in LAML), and regulators of apoptosis (such as LCL161 in OV and HNSC, and rTRAIL in BRCA), as well as chemotherapeutics (such as Bleomycin in BRCA, Temozolomide in COREAD and HNSC) (Figure 6C).

##### CELLector signatures are associated with differential prognosis

To further assess the clinical relevance of the CELLector signatures, we performed a systematic differential survival analysis across cancer types using publicly available survival data from cBioPortal (Cerami et al., 2012).

As for the drug association study presented in the previous section, comparing significance of top prognostic CELLector signatures with that of top prognostic individual CFEs showed that for STAD, LUAD and HNSC the former were more robustly associated with differential survival than the latters (Figure 6D, and Supplementary Figure S5D).

## Discussion

The translational potential of cancer preclinical studies is highly dependent on the clinical relevance of the employed *in vitro* models. Good models are required to capture the genomic heterogeneity of a cancer type under investigation and/or accurately represent alterations in relevant biological pathways.

We presented CELLector, a tool that allows scientists to select the most representative set of cell line models, maximising the covered genomic heterogeneity of the disease under consideration. The overall aim of the CELLector algorithm is to globally assess the quality of cancer *in vitro* models in terms of their ability to represent recurrent genomic subtypes detected in matched primary tumours, and to make available to the research community a user-controlled environment to perform such a task.

A key strength of CELLector is its generality: the algorithm can be applied to any disease for which *in vitro* models and matching primary/model genomic data are available. CELLector enables the systematic identification of recurrent tumour subtypes with paired genomic signatures, and selection of *in vitro* models based on the recurrence of these signatures. In addition, the algorithm identifies disease subtypes currently lacking representative models enabling prioritisation of new model development. To the best of our knowledge, CELLector represents the first computational method that ranks and selects cancer *in vitro* models, in a data driven way, across different cancer types, and without the need for expert knowledge about the primary disease under consideration. However, the model selection performed by CELLector can be flexibly tailored to fit the context of a study.

Here, we also demonstrated the power of CELLector analytical framework in bridging cancer patient genomics with cell line based pharmacogenomic studies. Our systematic analyses highlight the feasibility of linking the patient-defined CELLector signatures with differential drug response in cell lines, thus enabling *in-silico drug prescriptions* to precisely defined sub-cohorts of cancer patients, identifying (at the same time) sub-cohorts of patients that are not represented by any cell lines. This can be used for advising the generation of new *in vitro* models. CELLector analytical framework provides a powerful and clinically relevant way to associate tumour genotypes (together with their prevalence) with established or potential cancer therapies and supports the identification of complex and more robust markers of drug response.

Clinically relevant disease subtyping takes time and multiple resources. In recent years, an increasing number of studies have taken advantage of the availability of rich genomics/transcriptomics data for systematic molecular subclassification of tumours across tissues (Dawson et al., 2013; Guinney et al., 2015). Based on similar principles, CELLector can serve as a valuable tool to aid designing experimental studies minimising the risk of clinically relevant signal being missed due to ‘noise’ contributed by inclusion of less relevant models, or conversely identification of false positives due to strong signals from poor quality models. Addressing both of these issues will have direct and immediate implications on the quality of future *in vitro* experiments and in analysis of retrospective data derived from cancer cell lines.

## Supporting information

Suppl_Data_S1_TumourSubpopulationsRepresented-byAndLackingInVitroModels

Suppl_Data_S2_CELLector_Signature_BEMs_cell_lines

Suppl_Data_S3_ANOVA_inputs

Suppl_Data_S4_ANOVA_Results_across_13_cancer_types

Suppl_Data_S5_pvalue_comparisons_across_drugs

Suppl_Data_S6_CELLector_Signature_and_CFE_BEMs_primary_tumours

Suppl_Data_S7_Systematic_Differential_Survival_Analysis_Results

Supplemental_Information_User_Tutorial

## Author Contributions

H.N. contributed to algorithmic design, curated data, implemented, managed and documented the R package, worked on the R Shiny App development, performed systematic analyses, and wrote the manuscript; M.Y. implemented the first core function of the R package and performed systematic survival analysis; H.F., C.P contributed to manuscript editing/revising; E.S., J.S.R., M.J.G. revised the manuscript and contributed to the supervision of the study; F.I. conceived and designed the algorithm, implemented the R Shiny App, wrote the manuscript, and supervised the study. All authors read and approved the final manuscript.

## Acknowledgments

This work is funded by Open Targets project grant OTAR041. We thank Ian Dunham, Anneliese Speak, and Emanuel Gonçalves for critically reading the manuscript and providing useful feedback. We thank Fiona Behan, Patricia Jaaks and Evangelia Petsalaki for testing CELLector. We thank Patricia Jaaks, Lizzie Coker and Mathew Garnett for helping us mining/interpreting the signature/drug pharmacogenomic associations unveiled by our analysis. FI thanks Giorgia Iorio for her insightful comments on the visualisation methods used by our software.

## Supplementary Figures

**Supplementary Figure S1.**
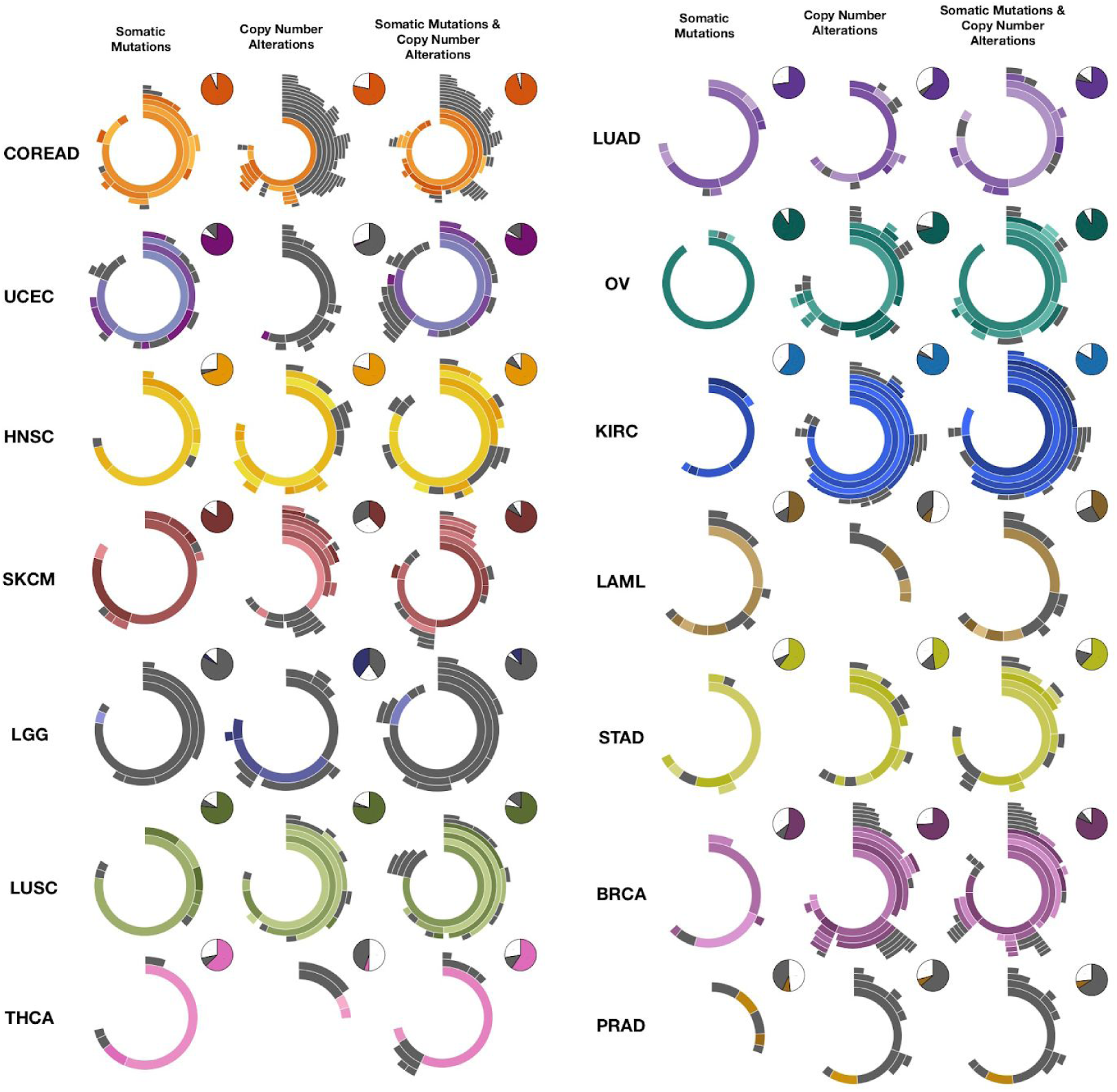
Systematic evaluation of tumour genomic heterogeneity and identification of tumour subtypes represented-by/lacking-representative *in vitro* models. The genomic landscape of 14 cancer types estimated using CELLector based on somatic mutation, copy number alterations and combination of both somatic mutations and copy number alterations in at least 2% of patient cohort. In each sunburst, segments represent tumour subtypes with those in grey indicating tumour subtypes lacking representative *in vitro* models. Pie charts represent proportion of patients that are not accounted for in the sunburst (white), or are accounted but not represented by any existing in vitro model (grey), or belonging to a subtype for which there is at least a representative in vitro model (color-coded respectively to cancer type).

**Supplementary Figure S2.**
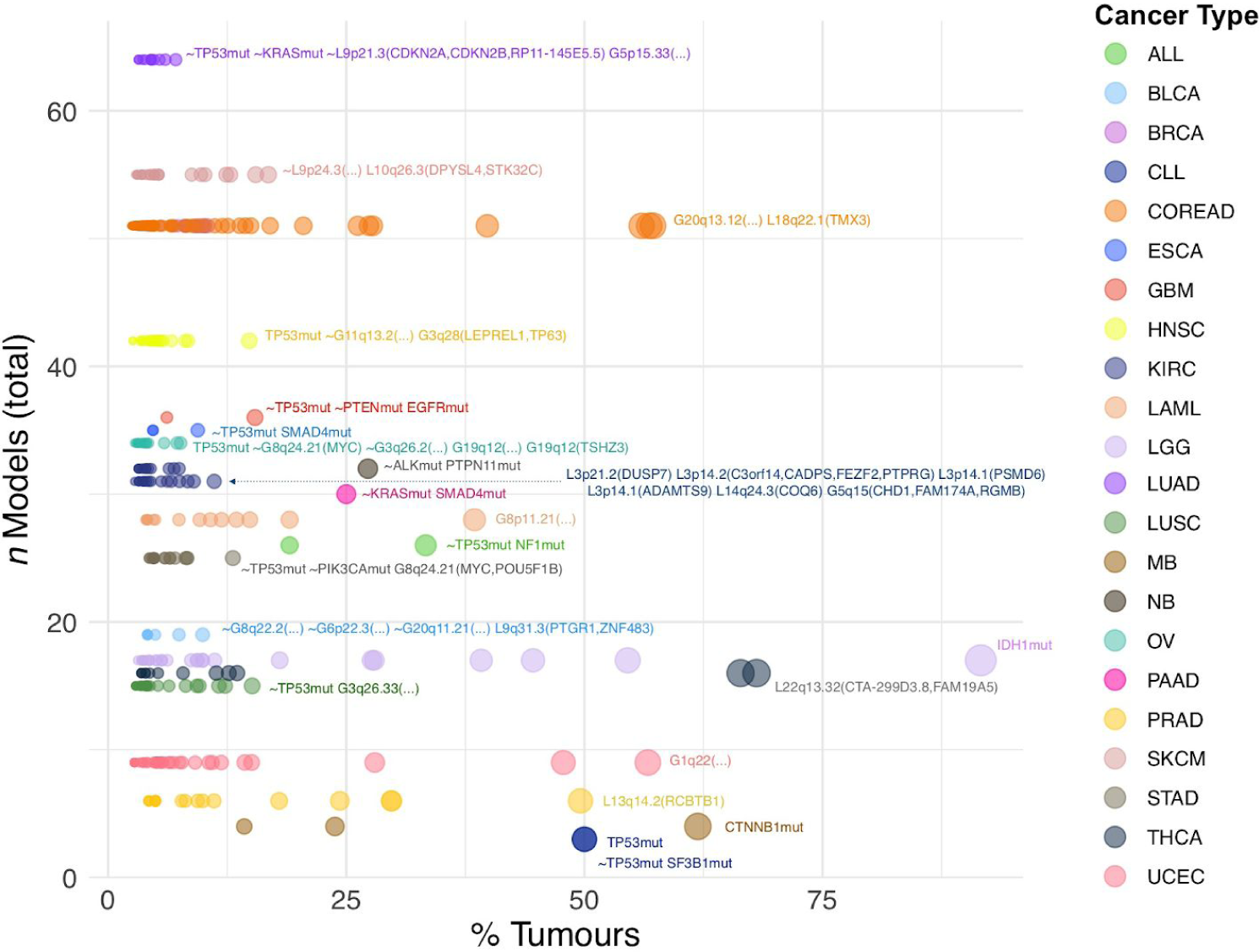
Patient subpopulations lacking representative *in vitro* models across 22 cancer types. Each point represents a patient subpopulation identified by CELLector (described by a signature) that does not have a representative cell line model when consider somatic mutations, copy number alteration and combination of both, occurring in at least 2% of patient cohort; Coordinates on the x-axis indicates the size of the subpopulation in terms of percentage of patients overall considered cohort. Coordinates on the y-axis indicate the total number of available tissue-matched cell line models. The CELLector signatures underlying the largest subpopulations are specified in the insets for each of the considered cancer types.

**Supplementary Figure S3.**
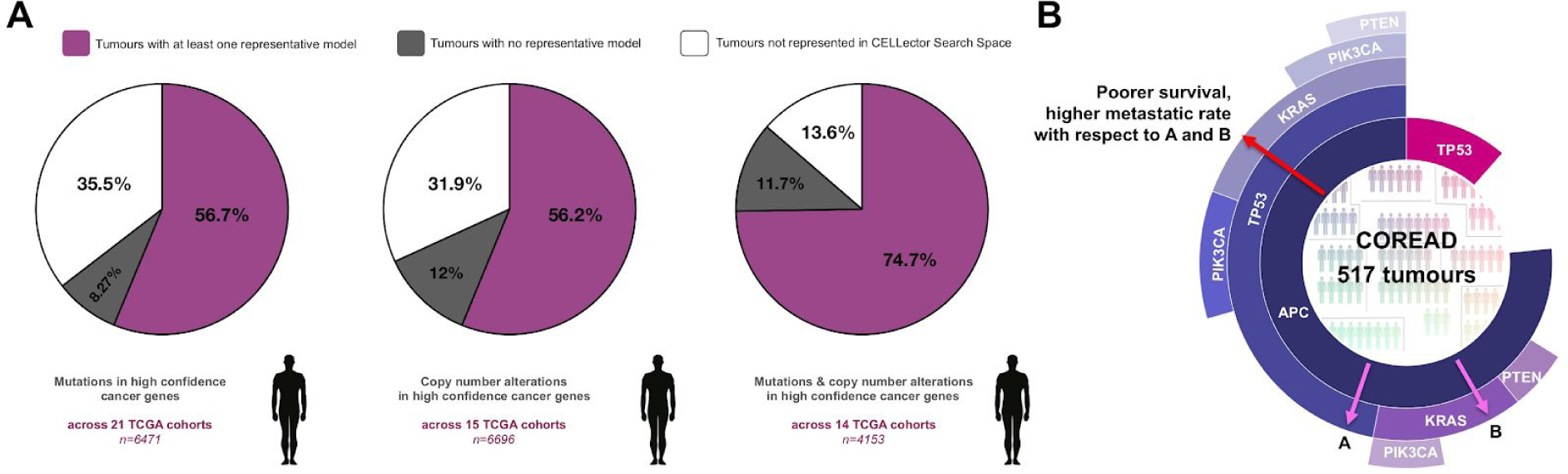
Patient represented-by/lacking suitable *in vitro* models, and clinical relevance of COREAD specific signatures. **A.** Ratios of patients with/without a representative *in vitro* model when pooling all cancer types. Each pie chart shows the proportion of tumours with no representative model (grey), with at least one representative model (purple) and that are not represented in CELLector search space (white). Different pie charts indicate CELLector analyses that consider i) somatic mutations, ii) copy number alterations and iii) combination of both somatic mutations and copy number alterations, respectively. **B.** A sunburst representation of COREAD patient subtypes identified by CELLector. Each segment represents a subtypes, with segment lengths indicating their recurrence. Concatenating the labels across nested segments provides the genomic signature describing the subtype in the outermost segment. Arrows indicate subtypes with differential prognosis.

**Supplementary Figure S4.**
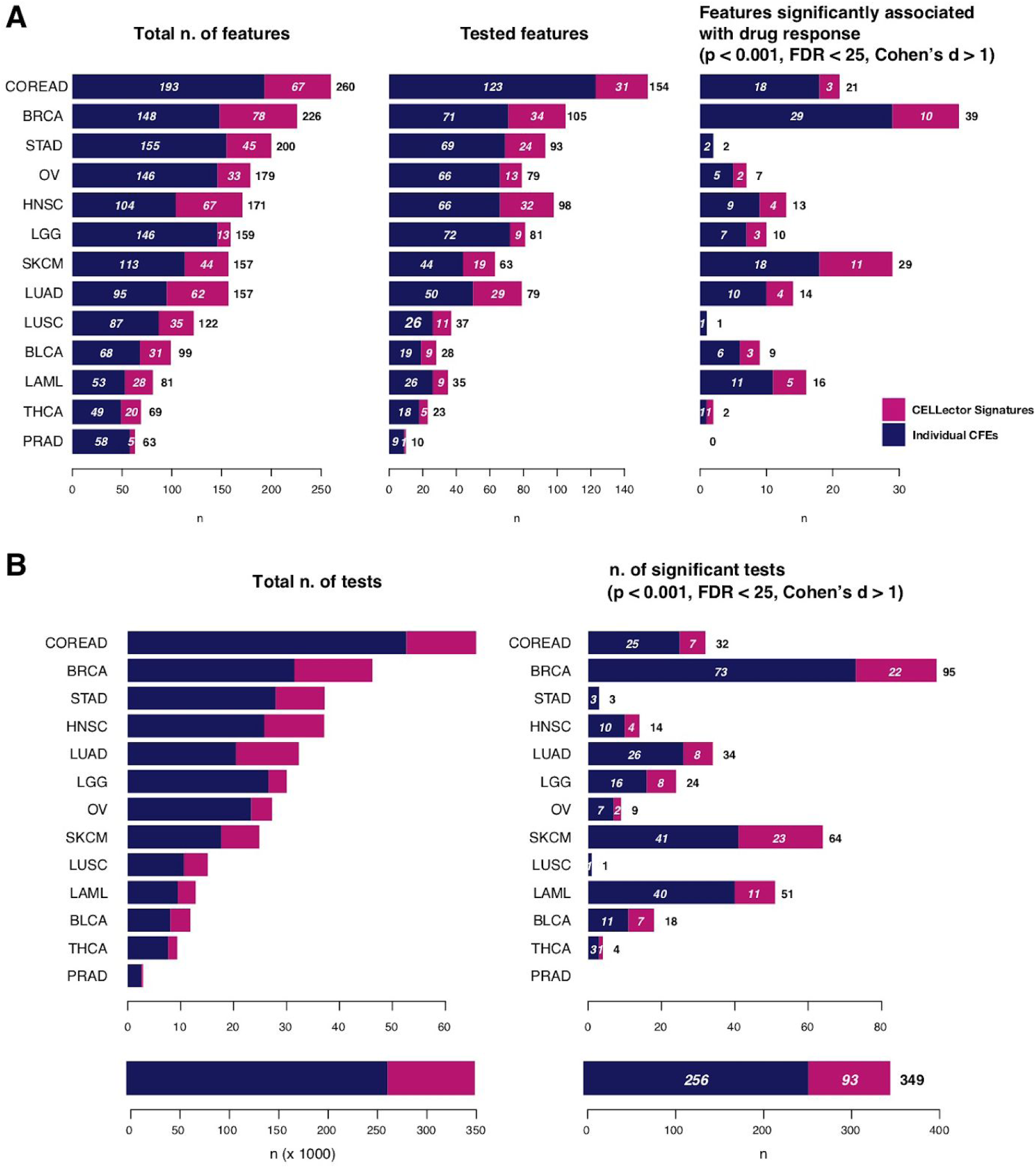
Summaries of systematic tissue specific Analysis of Variance (ANOVAs) across cancer types. **A.** (1) Total number of individual Cancer Functional Events (CFEs) and CELLector signatures. (ii) Number of CFEs and CELLector signatures used in the ANOVAs (occurring in at least 3 cell lines and less than *n* - 2 cell lines, with *n* = total number of cell lines for the cancer type under consideration). (iii) Number of CFEs and CELLector signatures involved in at least one statistically significant pharmacogenomic association. **B.** (i) Total number of tests performed in each cancer type specific analysis and grand total (bar in the bottom plot) and involving CFEs or CELLector signatures (as indicated by the different colours). (ii) Number of significant pharmacogenomic associations across cancer type specific ANOVAs and grand total (bar in the bottom plot) with colour scheme as for (i).

**Supplementary Figure S5.**
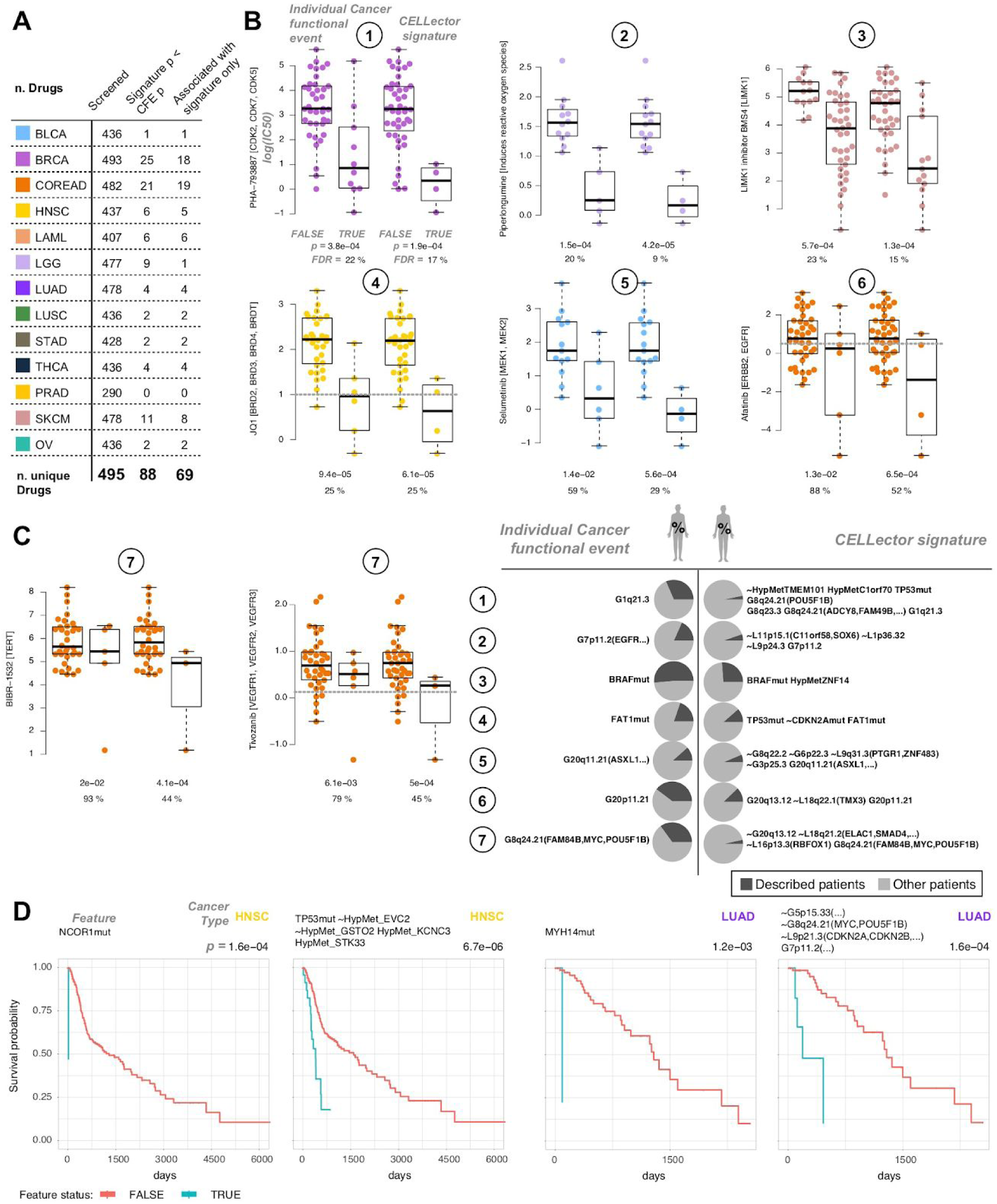
Systematic Analysis of Variance (ANOVA) and differential survival analysis outcomes. **A.** Number of drugs included in each cancer type specific ANOVA (1st column), for which there is at least one CELLector signatures associated with differential drug response more significantly than the top significant (p < 0.001) individual cancer functional event (CFE) (2nd column), for which there is at least one CELLector signature but no individual CFEs significantly (p < 0.001) associated with differential drug response (3rd column). **B.** Pairs of plots with examples of drug-signature associations (second plot in each pair) that are much more significant than the top significant drug-CFE association involving the same drug (first plot in each pair). Each circle represents a cell line with coordinate on the y-axis indicating the log IC50 of the drug specified in the y-axis label. In each individual plot, cell lines are partitioned into two groups based on the status of a genomic feature (TRUE or FALSE, indicating respectively the presence or absence of that feature). These features can be individual CFEs (first plot in each pair) or CELLector signatures (second plot in each pair), and are specified in the table in D. P-values and False Discovery Rates are from Analysis of Variance (ANOVA) tests assessing the extent of differential drug response across the considered dichotomies of cell lines. The boxes cover interquartile ranges with median lines drawn within them. Whiskers extend to a maximum of 1.5 times the size of the interquartile range. Point colors indicate the cancer type of origin of the cell lines. **C.** As for B but showing examples where CELLector signatures including MYC amplifications together with other CFEs (or their negation) are associated with drug response more significantly than MYC amplifications alone. **D.** Features (CFEs and CELLector signatures) whose status is used in the pair of plots in BC to dichotomised the cell lines and contrast their IC50s. The numerical labels on the left realise the mapping between each row and each pair of plots. Pie charts indicate percentage of patients harboring the indicated CFE (left panel) or CELLector signature (right panel), and whose cancer type matches that indicated by the color in the plot pairs. **E.** 2 Example Kaplan-Meier plots (respectively for HNSC and LUAD) showing CELLector signatures (2dn plot in each pair) predicting survival better that the top predictive individual CFE (1st plot in each pair). Shown Cox p-values are corrected by age and gender, and are significant at a FDR<25%.

## Supplementary Data

**Supplementary Data S1. Genomic signatures underlying tumour subpopulations represented-by and lacking *in vitro* models.**

**Supplementary Data S2. Status of patient-defined CELLector signatures in cell lines, across cancer types.**

**Supplementary Data S3. Patient-defined CELLector signatures and individual CFEs across cell lines used in systematic pharmacogenomic analysis.**

**Supplementary Data S4. Cancer-type specific ANOVA Results.**

**Supplementary Data S5. Comparison of CELLector Signature ANOVA p-values and individual CFE ANOVA p-values for all screened drugs across 13 cancer types.**

**Supplementary Data S6. CELLector signatures and individual CFEs across studied patient cohorts used in systematic survival analysis.**

**Supplementary Data S7. Systematic differential survival analysis results.**

## STAR Methods

### KEY RESOURCES TABLE

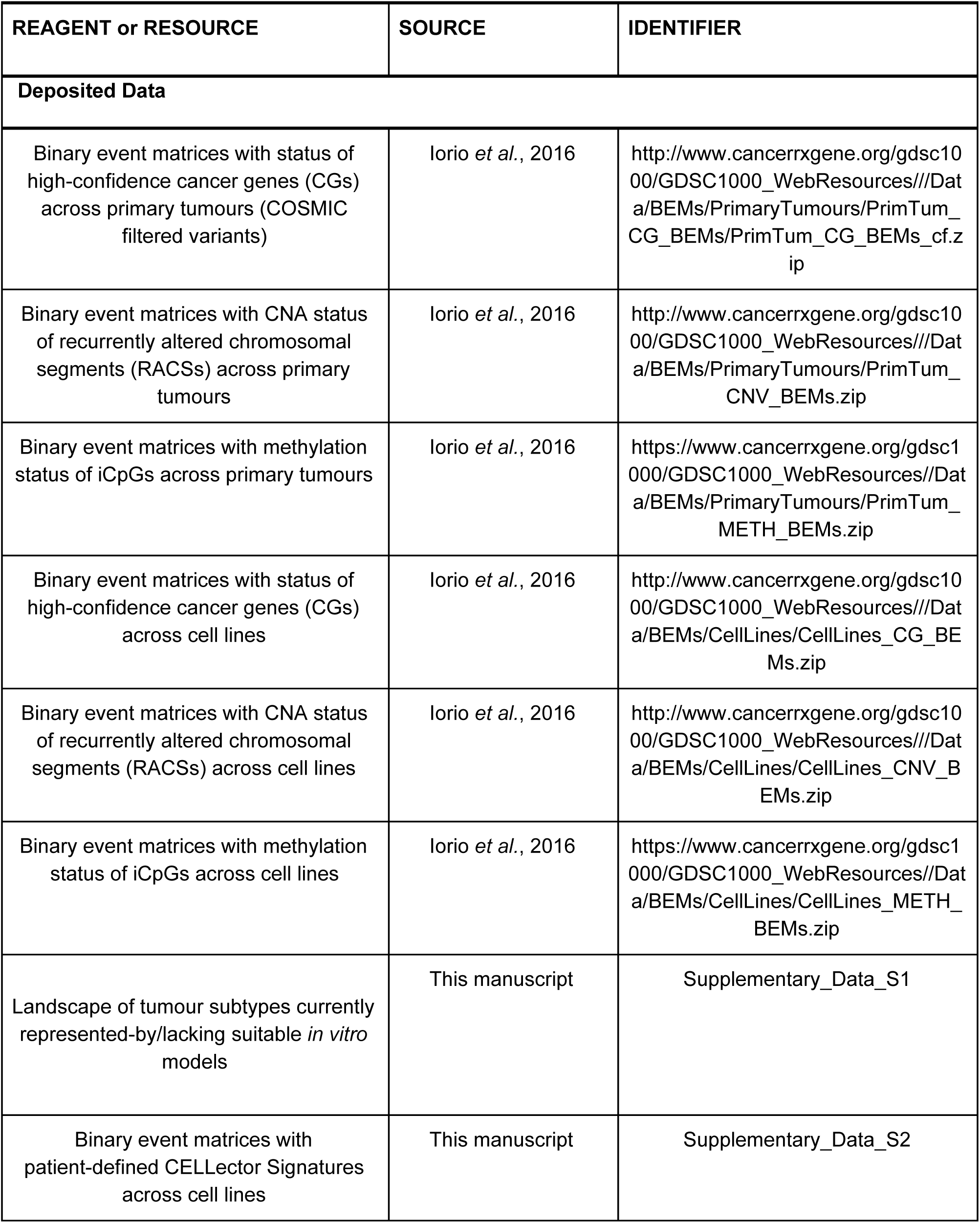
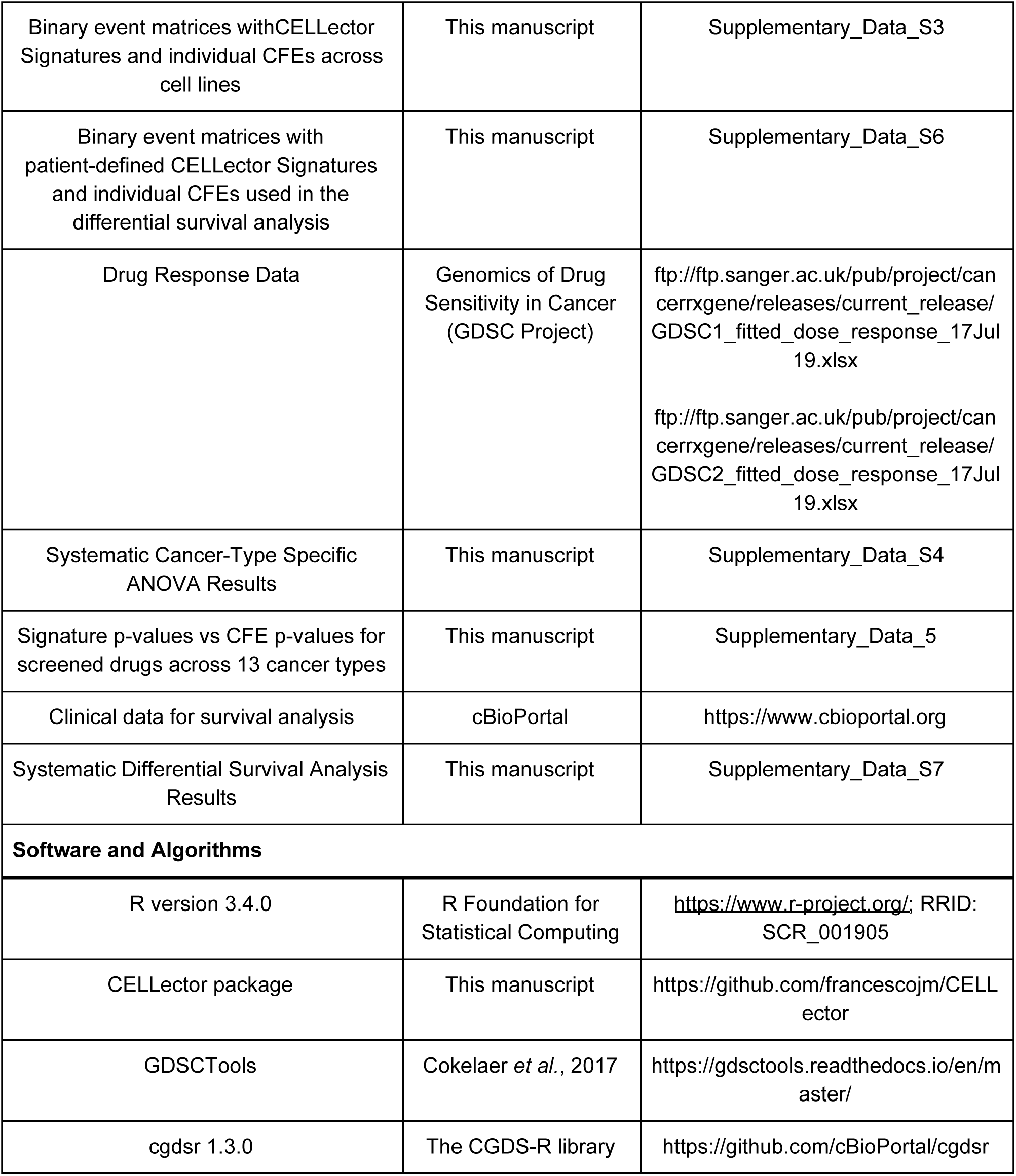

### CONTACT FOR REAGENT AND RESOURCE SHARING

Further information and requests for resources and reagents should be directed to and will be fulfilled by the Lead Contact, Francesco Iorio.

### METHOD DETAILS

#### Implementation

The CELLector algorithm and interactive visualisation tools are implemented in R and available as an open-source R package (code available at https://github.com/francescojm/CELLector, interactive vignette available at http://rpubs.com/francescojm/CELLector, user manual available at https://github.com/francescojm/CELLector/blob/master/CELLector.pdf) and R Shiny web application (deployed at https://ot-cellector.shinyapps.io/cellector_app/, code available at https://github.com/francescojm/CELLector_App).

#### Genomics data

CELLector provides built-in genomics data for disease-matched primary tumours and cell lines derived from 16 cancer types, encompassing the characterisation of 4,550 tumours and 499 immortalised and commercially available cancer cell lines (Supplemental Information Table S1), and accounting for somatic mutations, copy number alterations and hypermethylated gene promoters for high-confidence cancer genes and recurrently altered chromosomal segments, i.e. cancer functional events (CFEs). These CFEs are described in Iorio *et al*., 2016 and corresponding data were obtained from the accompanied web-portal (http://www.cancerrxgene.org/gdsc1000/).

#### QUANTIFICATION AND STATISTICAL ANALYSIS: The CELLector algorithm

In the analytical framework of CELLector, the genomic background of a cohort of patients affected by a given cancer type is represented as a binary tree whose topology is determined by the most-frequently observed combinations of molecular alterations (item-sets) and their supports, i.e. the fraction of patients in which these alterations occur simultaneously. This tree is built recursively by sequential applications of the *Eclat* algorithm (Zaki et al., 1997) as follows. The tree construction starts from the root, modelling the combination of genomic alterations (item-set) with the largest support across the entire cohort. Then two sibling nodes are included, modelling the item-sets with the greatest support when considering the population supporting the item-set of the parent node (right sibling node) and its complementary population (left sibling node). This is recursively performed at each new node included in the tree if the corresponding modelled item-set is supported by at least a user-defined ratio of patients in the considered patient subpopulation (for example 5%).

Subsequently, a logic AND formula *F* is assigned to each node *x*, considering the path to *x* from the root of the three. For each node *n* on this path (including the terminal ones) the corresponding modelled item-set is added to *F* as a term, negated if *n* is a left sibling (complement) node. Finally, a given cell line in the built-in collection is mapped to a node *n* if its genomic background satisfies *F(n)*.

The algorithm continues with a guided deep-first-visit of the obtained tree, which return all the identified subtypes as a sorted list, as detailed in the following pseudo code:

#### Variables and initial settings

*Q* = an empty queue

*T* = a CELLector searching space

*r* = the root of T

*U* = a set of nodes that have not been visited yet

*Idx* = a queue index *CurrentNode* = *r*

*Idx* = −1

*U* = all the nodes of T

#### Algorithm

~~~
While *U* is not empty
     remove CurrentNode from U
     While *CurrentNode* as a left child
          Add *CurrentNode* to the queue
          CurrentNode = left child of CurrentNode

end

Add the right children of all the nodes in *Q* to *Q* (by level) and remove them from *U*
If there are right nodes in *Q* in position > *Idx* then
         Advance *Idx* to the first right node in *Q* in a position > *Idx
         CurrentNode* = *Q[Idx]*
end
Return *Q*
~~~

Finally, a Cell Line map is built by considering all subtypes (nodes) as they appear in *Q*, with corresponding signatures and mapped cell lines.

*N* Cell lines are selected among those appearing in the first *N* entries of this map with a heuristic method, minimizing the number of nodes each selected cell line is mapped onto.

## DATA AND SOFTWARE AVAILABILITY

Source code for the R package and the R Shiny app is publicly available on GitHub: Package code https://github.com/francescojm/CELLector

Package interactive vignette http://rpubs.com/francescojm/CELLector

Package user manual https://github.com/francescojm/CELLector/blob/master/CELLector.pdf, App deployed at https://ot-cellector.shinyapps.io/cellector_app/

App code available at https://github.com/francescojm/CELLector_App.

Detailed instructions on how to install the R package, run the CELLector analysis and interactively explore the results are also provided in the GitHub repository and in the Supplemental Information, together with a tutorial with instructions how to set up the analysis, interactively explore the results and execute CELLector on example case studies.

### Pharmacogenomic analysis

Patient-defined CELLector signatures for the systematic drug association studies were obtained by running 6 different CELLector analyses that considered either i) mutations only (M), ii) copy number alterations only (C), iii) mutations and copy number alterations (MC), iv) mutations and methylation (MH), v) copy number alterations and methylation (CH) or vi) mutations, copy number alterations and methylation (MCH) prevalent in at least 2% of analysed patient cohort across 13 cancer types (LUAD, COREAD, HNSC, BRCA, PRAD, SKCM, BLCA, STAD, LUSC, THCA, LGG, LAML, OV). The obtained patient-defined signatures were then mapped onto tissue-matched cell lines (Supplementary Data S2) and used together with individual Cancer Functional Events (CFEs, from (Iorio et al., 2016)) in systematic cancer-specific ANOVAs (Supplementary Data S3) to test if they were statistically associated with differential response to 495 compounds (Picco et al., 2019). The patient-defined signatures from different CELLector analyses and individual CFEs were collated into one input feature matrix per cancer type (Supplementary Figure S4). Signature duplications were discarded (e.g. the same signature was present in multiple CELLector analyses or signature and CFE were the same, Supplementary Data S3). Only signatures or CFEs with at least 3 mapped cell lines and less than *n* - 2 mapped cell lines (where *n* is the total number of available cell lines for the tissue under consideration, Supplemental Information Table S1) where included in the ANOVAs (Supplementary Data S3). Statistical details on the ANOVAs are provided in (Iorio et al., 2016). Particularly, we tested associations between pattern of drug IC50s and the presence of individual CFEs or CELLector signatures including the microsatellite instability (MSI) status of the cell lines as a cofactor (for cancer types where at least 3 MSI instable cell lines were present). A total of 13 analyses (referred for simplicity as cancer-type-specific ANOVAs in the main text and below) were performed. Each ANOVA was performed using the analytical framework implemented in the GDSCtools Python package (Cokelaer et al., 2018) (https://github.com/CancerRxGene/gdsctools). For all tested drug-CFE/signature associations, effect size estimations versus pooled s.d. (quantified using Cohen’s d), effect sizes versus individual s.d. (quantified using two different Glass’s Δ metrics, for the positive and the negative populations separately), p values and all other statistical scores were obtained from the fitted models. *p-*values from the same cancer type specific ANOVA were corrected all together using the Benjamini-Hochberg. Results from all the tests are reported in Supplementary Data S4.

### Survival analysis

Patient-defined CELLector signatures for the systematic survival analysis were obtained by running 3 different CELLector analyses that considered either i) mutations and methylation (MH), ii) copy number alterations and methylation (CH) or iii) mutations, copy number alterations and methylation (MCH) prevalent in at least 2% of analysed patient cohort across 13 cancer types (LUAD, COREAD, HNSC, BRCA, PRAD, SKCM, BLCA, STAD, LUSC, THCA, LGG, LAML, OV; Supplementary Data S6). CELLector signatures from mutations only (M), copy number alterations only and mutations and copy number alterations (MC) were not included in this analysis due to asymmetric patient identifiers. The obtained patient-defined signatures and individual Cancer Functional Events (CFEs, from (Iorio et al., 2016)) were collated into one feature matrice per cancer type (Supplementary Data S6) and used in systematic differential survival analysis. In cases where patient-defined signatures were duplicated (e.g. the same signature was present in multiple CELLector analyses or signature and CFE were the same), the prevalence of that signature in the studied patient cohorts was considered; the duplicate with higher number of underlining patients was retained. The clinical data for systematic survival analysis were downloaded from cBioPortal using the cgdsr 1.3.0 R package. The Cox *P*-values for either signatures or CFEs were corrected by age and gender (Supplementary Data S7).

### Supplemental Information

Supplemental Information includes one table, two case studies, instructions how to install and run different CELLector modalities (e.g. R package, online and local R Shiny App) and user tutorials demonstrating the full functionality of CELLector app.

## References

Ahmed, D., Eide, P.W., Eilertsen, I.A., Danielsen, S.A., Eknæs, M., Hektoen, M., Lind, G.E., and Lothe, R.A. (2013). Epigenetic and genetic features of 24 colon cancer cell lines. Oncogenesis 2, e71–e71.

Barretina, J., Caponigro, G., Stransky, N., Venkatesan, K., Margolin, A.A., Kim, S., Wilson, C.J., Lehár, J., Kryukov, G.V., Sonkin, D., et al. (2012). The Cancer Cell Line Encyclopedia enables predictive modelling of anticancer drug sensitivity. Nature 483, 603–607.

Beaufort, C.M., Helmijr, J.C.A., Piskorz, A.M., Hoogstraat, M., Ruigrok-Ritstier, K., Besselink, N., Murtaza, M., van Ĳcken, W.F.J., Heine, A.A.J., Smid, M., et al. (2014). Ovarian cancer cell line panel (OCCP): clinical importance of in vitro morphological subtypes. PLoS One 9, e103988.

Behan, F.M., Iorio, F., Picco, G., Gonçalves, E., Beaver, C.M., Migliardi, G., Santos, R., Rao, Y., Sassi, F., Pinnelli, M., et al. (2019). Prioritization of cancer therapeutic targets using CRISPR-Cas9 screens. Nature 568, 511–516.

Cerami, E., Gao, J., Dogrusoz, U., Gross, B.E., Sumer, S.O., Aksoy, B.A., Jacobsen, A., Byrne, C.J., Heuer, M.L., Larsson, E., et al. (2012). The cBio cancer genomics portal: an open platform for exploring multidimensional cancer genomics data. Cancer Discov. 2, 401–404.

Cokelaer, T., Chen, E., Iorio, F., Menden, M.P., Lightfoot, H., Saez-Rodriguez, J., and Garnett, M.J. (2018). GDSCTools for mining pharmacogenomic interactions in cancer. Bioinformatics 34, 1226–1228.

Domcke, S., Sinha, R., Levine, D.A., Sander, C., and Schultz, N. (2013). Evaluating cell lines as tumour models by comparison of genomic profiles. Nat. Commun. 4, 2126.

van Dyk, E., Reinders, M.J.T., and Wessels, L.F.A. (2013). A scale-space method for detecting recurrent DNA copy number changes with analytical false discovery rate control. Nucleic Acids Res. 41, e100.

Forbes, S.A., Beare, D., Boutselakis, H., Bamford, S., Bindal, N., Tate, J., Cole, C.G., Ward, S., Dawson, E., Ponting, L., et al. (2017). COSMIC: somatic cancer genetics at high-resolution. Nucleic Acids Res. 45, D777–D783.

Garnett, M.J., Edelman, E.J., Heidorn, S.J., Greenman, C.D., Dastur, A., Lau, K.W., Greninger, P., Thompson, I.R., Luo, X., Soares, J., et al. (2012). Systematic identification of genomic markers of drug sensitivity in cancer cells. Nature 483, 570–575.

Gonzalez-Perez, A., Perez-Llamas, C., Deu-Pons, J., Tamborero, D., Schroeder, M.P., Jene-Sanz, A., Santos, A., and Lopez-Bigas, N. (2013). IntOGen-mutations identifies cancer drivers across tumor types. Nat. Methods 10, 1081–1082.

Gundem, G., Perez-Llamas, C., Jene-Sanz, A., Kedzierska, A., Islam, A., Deu-Pons, J., Furney, S.J., and Lopez-Bigas, N. (2010). IntOGen: integration and data mining of multidimensional oncogenomic data. Nat. Methods 7, 92–93.

Han, J., Pei, J., and Kamber, M. (2011). Data Mining: Concepts and Techniques (Elsevier).

Hodis, E., Watson, I.R., Kryukov, G.V., Arold, S.T., Imielinski, M., Theurillat, J.-P., Nickerson, E., Auclair, D., Li, L., Place, C., et al. (2012). A landscape of driver mutations in melanoma. Cell 150, 251–263.

Ince, T.A., Sousa, A.D., Jones, M.A., Harrell, J.C., Agoston, E.S., Krohn, M., Selfors, L.M., Liu, W., Chen, K., Yong, M., et al. (2015). Characterization of twenty-five ovarian tumour cell lines that phenocopy primary tumours. Nat. Commun. 6, 7419.

Iorio, F., Knijnenburg, T.A., Vis, D.J., Bignell, G.R., Menden, M.P., Schubert, M., Aben, N., Gonçalves, E., Barthorpe, S., Lightfoot, H., et al. (2016). A Landscape of Pharmacogenomic Interactions in Cancer. Cell 166, 740–754.

Jiang, G., Zhang, S., Yazdanparast, A., Li, M., Pawar, A.V., Liu, Y., Inavolu, S.M., and Cheng, L. (2016). Comprehensive comparison of molecular portraits between cell lines and tumors in breast cancer. BMC Genomics 17 *Suppl 7*, 525.

Kaur, M. (2014). ECLAT Algorithm for Frequent Itemsets Generation. International Journal of Computer Systems 1, 82–84.

Kentsis, A., Reed, C., Rice, K.L., Sanda, T., Rodig, S.J., Tholouli, E., Christie, A., Valk, P.J.M., Delwel, R., Ngo, V., et al. (2012). Autocrine activation of the MET receptor tyrosine kinase in acute myeloid leukemia. Nat. Med. 18, 1118–1122.

Kojima, K., Konopleva, M., McQueen, T., O’Brien, S., Plunkett, W., and Andreeff, M. (2006). Mdm2 inhibitor Nutlin-3a induces p53-mediated apoptosis by transcription-dependent and transcription-independent mechanisms and may overcome Atm-mediated resistance to fludarabine in chronic lymphocytic leukemia. Blood 108, 993–1000.

Medico, E., Russo, M., Picco, G., Cancelliere, C., Valtorta, E., Corti, G., Buscarino, M., Isella, C., Lamba, S., Martinoglio, B., et al. (2015). The molecular landscape of colorectal cancer cell lines unveils clinically actionable kinase targets. Nature Communications 6.

van der Meer, D., Barthorpe, S., Yang, W., Lightfoot, H., Hall, C., Gilbert, J., Francies, H.E., and Garnett, M.J. (2019). Cell Model Passports—a hub for clinical, genetic and functional datasets of preclinical cancer models. Nucleic Acids Res. 47, D923–D929.

Mouradov, D., Sloggett, C., Jorissen, R.N., Love, C.G., Li, S., Burgess, A.W., Arango, D., Strausberg, R.L., Buchanan, D., Wormald, S., et al. (2014). Colorectal cancer cell lines are representative models of the main molecular subtypes of primary cancer. Cancer Res. 74, 3238–3247.

Picco, G., Chen, E.D., Alonso, L.G., Behan, F.M., Gonçalves, E., Bignell, G., Matchan, A., Fu, B., Banerjee, R., Anderson, E., et al. (2019). Functional linkage of gene fusions to cancer cell fitness assessed by pharmacological and CRISPR-Cas9 screening. Nat. Commun. 10, 2198.

Qiu, Z., Zou, K., Zhuang, L., Qin, J., Li, H., Li, C., Zhang, Z., Chen, X., Cen, J., Meng, Z., et al. (2016). Hepatocellular carcinoma cell lines retain the genomic and transcriptomic landscapes of primary human cancers. Sci. Rep. 6, 27411.

Sanz-Garcia, E., Argiles, G., Elez, E., and Tabernero, J. (2017). BRAF mutant colorectal cancer: prognosis, treatment, and new perspectives. Ann. Oncol. 28, 2648–2657.

Schell, M.J., Yang, M., Teer, J.K., Lo, F.Y., Madan, A., Coppola, D., Monteiro, A.N.A., Nebozhyn, M.V., Yue, B., Loboda, A., et al. (2016). A multigene mutation classification of 468 colorectal cancers reveals a prognostic role for APC. Nat. Commun. 7, 11743.

Shoemaker, R.H. (2006). The NCI60 human tumour cell line anticancer drug screen. Nat. Rev. Cancer 6, 813–823.

Sinha, R., Schultz, N., and Sander, C. (2015). Comparing cancer cell lines and tumor samples by genomic profiles. bioRxiv.

Sinha, R., Winer, A.G., Chevinsky, M., Jakubowski, C., Chen, Y.-B., Dong, Y., Tickoo, S.K., Reuter, V.E., Russo, P., Coleman, J.A., et al. (2017). Analysis of renal cancer cell lines from two major resources enables genomics-guided cell line selection. Nat. Commun. 8, 15165.

Storey, J.D., and Tibshirani, R. (2003). Statistical significance for genomewide studies. Proc. Natl. Acad. Sci. U. S. A. 100, 9440–9445.

Sun, Y., and Liu, Q. (2015). Deciphering the Correlation between Breast Tumor Samples and Cell Lines by Integrating Copy Number Changes and Gene Expression Profiles. Biomed Res. Int. 2015, 901303.

Tsherniak, A., Vazquez, F., Montgomery, P.G., Weir, B.A., Kryukov, G., Cowley, G.S., Gill, S., Harrington, W.F., Pantel, S., Krill-Burger, J.M., et al. (2017). Defining a Cancer Dependency Map. Cell 170, 564–576.e16.

Vincent, K.M., Findlay, S.D., and Postovit, L.M. (2015). Assessing breast cancer cell lines as tumour models by comparison of mRNA expression profiles. Breast Cancer Research 17.

Yang, W., Soares, J., Greninger, P., Edelman, E.J., Lightfoot, H., Forbes, S., Bindal, N., Beare, D., Smith, J.A., Richard Thompson, I., et al. (2012). Genomics of Drug Sensitivity in Cancer (GDSC): a resource for therapeutic biomarker discovery in cancer cells. Nucleic Acids Research 41, D955–D961.

Zhang, J., Baran, J., Cros, A., Guberman, J.M., Haider, S., Hsu, J., Liang, Y., Rivkin, E., Wang, J., Whitty, B., et al. (2011). International Cancer Genome Consortium Data Portal—a one-stop shop for cancer genomics data. Database 2011.

Zhao, N., Liu, Y., Wei, Y., Yan, Z., Zhang, Q., Wu, C., Chang, Z., and Xu, Y. (2017). Optimization of cell lines as tumour models by integrating multi-omics data. Brief. Bioinform. 18, 515–529.

