## Supplemental_Information_User_Tutorial for "CELLector: Genomics Guided Selection of Cancer *in vitro* Models"

[8] Lead Contact

### 1. Supplementary Table

| **TCGA Label** | **Cancer Type** | **n. of primary tumours** | **n. of Cell lines** |
|---|---|---|---|
| GBM | Glioblastoma Multiforme | 302 | 36 |
| LUAD | Lung Adenocarcinoma | 183 | 64 |
| KIRC | Kidney Renal Clear Cell Carcinoma | 451 | 31 |
| COREAD | Colon Adenocarcinoma and Rectum Adenocarcinoma | 517 | 51 |
| HNSC | Head and Neck Squamous Cell Carcinoma | 299 | 42 |
| BRCA | Breast Invasive Carcinoma | 759 | 51 |
| PRAD | Prostate Adenocarcinoma | 179 | 6 |
| SKCM | Skin Cutaneous Melanoma | 251 | 55 |
| BLCA | Bladder Urothelial Carcinoma | 95 | 19 |
| STAD | Stomach Adenocarcinoma | 107 | 25 |
| LUSC | Lung Squamous Cell Carcinoma | 186 | 15 |
| UCEC | Uterine Corpus Endometrial Carcinoma | 228 | 9 |
| THCA | Thyroid Carcinoma | 321 | 16 |
| LGG | Brain Lower Grade Glioma | 169 | 17 |
| LAML | Acute Myeloid Leukemia | 186 | 28 |
| OV | Ovarian Serous Cystadenocarcinoma | 317 | 34 |
| | | 4550 | 499 |

**Table S1 - Summary of CELLector built-in primary tumour and cell line genomics data sets.** CELLector makes use of somatic mutations, copy number alterations and methylation data for primary tumours and cell lines derived from 16 different tissue types, enabling genomics-guided evaluation of a total of 499 cancer *in vitro* models. These data are curated as described in *Iorio et al., Cell 2016*.

### 2. CELLector usage modalities

CELLector can be used in three different modalities:

- (i) as an R package (R code available at: https://github.com/francescojm/CELLector);
- (ii) as an online R shiny App (deployed at: https://ot-cellector.shinyapps.io/CELLector_App/);
- (iii) as an R shiny App locally, within Rstudio, code available at: https://github.com/francescojm/CELLector_App).

### 3. CELLector R package

A user manual with extensive documentation is available at: https://github.com/francescojm/CELLector/blob/master/CELLector.pdf and an interactive vignette with instructions to perform a simple quick analysis is available at: http://rpubs.com/francescojm/CELLector. A static version of this vignette is reported below.

#### Package installation

The R package is available on github at https://github.com/francescojm/CELLector. We recommend using it within Rstudio (https://www.rstudio.com/). To install it the following commands should be executed:

```
library(devtools)
install_github("Francescojm/CELLector")
library(CELLector)
```

This will install the following additional libraries

```
arules, dplyr, stringr, data.tree, sunburstR, igraph, collapsibleTree, methods
```

all publicly available on CRAN or Bioconductor. The package comes with built-in data objects containing genomic data for large cohorts of primary tumours and cancer cell lines (from Iorio *et al*, Cell 2016).

#### Selecting Colorectal Cancer Cell Lines that are representative of a defined patients' sub-population

In this simple case study scenario, we want to select the 5 most clinically relevant *in vitro* models that best represent the genomic diversity of TP53 mutated colorectal tumours. The models we wish to select should be microsatellite stable and harbour at least one alteration in the following signalling pathways: *PI3K-AKT-MTOR signalling*, *RAS-RAF-MEK-ERK/JNK signalling* and *WNT signalling*. Finally, we want the model selection to be guided based on somatic mutations and copy number alterations that are observed in at least 3% of the studied TP53 mutant tumour cohort.

After loading the package, we need to load the following data objects

```
library(CELLector)

## Somatic mutations and copy number alterations found in primary tumours and cell

## lines

data(CELLector.PrimTum.BEMs)

data(CELLector.CellLine.BEMs)

## Sets of Cancer Functional Events (CFEs: somatic mutations and copy number

## alterations) involving genes in predefined key cancer pathways

data(CELLector.Pathway_CFEs)

## Objects used for decoding cna CFEs identifiers
```

```
data(CELLector.CFEs.CNAid_mapping)

data(CELLector.CFEs.CNAid_decode)
```

Subsequently we specify the Colon/Rectal Adenocarcinoma (COREAD) TCGA label to select the cancer type to analyse. Other available cancer types in this version of the package are: *BLCA, BRCA, COREAD, GBM, HNSC, KIRC, LAML, LGG, LUAD, LUSC, OV, PRAD, SKCM, STAD, THCA, UCEC.* Finally, we remove possible sample identifier duplications from the primary tumour dataset.

```
### Change the following two lines to work with a different cancer type

tumours_BEM<-CELLector.PrimTum.BEMs$COREAD

CELLlineData<-CELLector.CellLine.BEMs$COREAD

### unicize the sample identifiers for the tumour data

tumours_BEM<-CELLector.unicizeSamples(tumours_BEM)
```

At this point, we are ready to build the CELLector searching space, which will contain the most recurrent patients subtypes with matched signatures of CFEs. The value of the arguments of the below function specify that we want to look only at TP53 mutant cancer patients, with alterations in genes belonging to three different cancer pathways.

```
### building a CELLector searching space focusing on three pathways

### and TP53 mutatant patients only

CSS<-CELLector.Build_Search_Space(ctumours = t(tumours_BEM), verbose = FALSE,
minGlobSupp = 0.03, cancerType = 'COREAD', pathwayFocused = c("RAS-RAF-MEK-ERK / JNK
signaling", "PI3K-AKT-MTOR signaling", "WNT signaling"),

pathway_CFEs = CELLector.Pathway_CFEs, cnaIdMap = CELLector.CFEs.CNAid_mapping,
cnaIdDecode = CELLector.CFEs.CNAid_decode, subCohortDefinition='TP53')
```

The searching space is stored in a list, whose element can be inspected as follows:

```
### visualising the CELLector searching space as a binary tree
CSS$TreeRoot
```

```
#>                          levelName
#> 1  1 APCmut
#> 2   ¦--2 KRASmut
#> 3   ¦   ¦--14 L17p12
#> 4   ¦   °--15 L17p12
#> 5   °--3 L17p12
#> 6       ¦--4 PIK3CAmut
#> 7       ¦   ¦--10 KRASmut
#> 8       ¦   °--11 PTENmut
#> 9       ¦       °--12 KRASmut
#> 10      ¦           °--13 NF1mut
#> 11      °--5 KRASmut
#> 12          ¦--6 PTENmut
#> 13          ¦   ¦--8 L5q21.1
#> 14          ¦   °--9 PIK3CAmut
#> 15          °--7 PIK3CAmut
```

```
### visualising the first attributes of the tree nodes CSS$navTable[,1:11]
```

```
#>    Idx  Items                      ItemsDecoded        Type Parent.Idx
#> 1    1    APC                            APCmut        root          0
#> 2    2   KRAS                           KRASmut Right.Child          1
#> 3    3  cna27 L17p12(DNAH9,MAP2K4,SHISA6,ZNF18)  Left.Child          1
#> 4    4 PIK3CA                         PIK3CAmut Right.Child          3
#> 5    5   KRAS                           KRASmut  Left.Child          3
#> 6    6   PTEN                           PTENmut Right.Child          5
#> 7    7 PIK3CA                         PIK3CAmut  Left.Child          5
#> 8    8  cna63                      L5q21.1(...) Right.Child          6
#> 9    9 PIK3CA                         PIK3CAmut  Left.Child          6
#> 10  10   KRAS                           KRASmut Right.Child          4
#> 11  11   PTEN                           PTENmut  Left.Child          4
#> 12  12   KRAS                           KRASmut  Left.Child         11
#> 13  13    NF1                            NF1mut  Left.Child         12
#> 14  14  cna27 L17p12(DNAH9,MAP2K4,SHISA6,ZNF18) Right.Child          2
#> 15  15  cna27 L17p12(DNAH9,MAP2K4,SHISA6,ZNF18)  Left.Child          2
#>    AbsSupport CurrentTotal PercSupport GlobalSupport Left.Child.Index
#> 1         243          302   0.8046358    0.80463576                3
#> 2          26           59   0.4406780    0.08609272               15
#> 3         131          243   0.5390947    0.43377483                5
#> 4          52          112   0.4642857    0.17218543               11
#> 5          54          131   0.4122137    0.17880795                7
#> 6          26           77   0.3376623    0.08609272                9
#> 7          22           54   0.4074074    0.07284768                0
#> 8          12           51   0.2352941    0.03973510                0
#> 9          15           26   0.5769231    0.04966887                0
#> 10         22           60   0.3666667    0.07284768                0
#> 11         28           52   0.5384615    0.09271523               12
#> 12         18           28   0.6428571    0.05960265               13
#> 13         11           18   0.6111111    0.03642384                0
#> 14         12           33   0.3636364    0.03973510                0
#> 15         14           26   0.5384615    0.04635762                0
#>    Right.Child.Index
#> 1                  2
#> 2                 14
#> 3                  4
#> 4                 10
#> 5                  6
#> 6                  8
#> 7                  0
#> 8                  0
#> 9                  0
#> 10                 0
#> 11                 0
#> 12                 0
#> 13                 0
#> 14                 0
#> 15                 0
```

```r
### visualising the sub-cohort of patients whose genome satisfies the rule of the 4th
### node

str_split(CSS$navTable$positivePoints[4],',')
```

```
#> [[1]]
#>  [1] "TCGA-A6-5662_29"  "TCGA-A6-6649_38"  "TCGA-A6-6654_42"
#>  [4] "TCGA-AA-3663_106" "TCGA-AA-3713_126" "TCGA-AA-3977_174"
#>  [7] "TCGA-AD-6899_228" "TCGA-AD-6963_230" "TCGA-AF-2687_233"
#> [10] "TCGA-AG-3731_272" "TCGA-AG-4015_295" "TCGA-AH-6544_316"
#> [13] "TCGA-AH-6643_318" "TCGA-AU-3779_320" "TCGA-AY-6197_326"
#> [16] "TCGA-AZ-4616_330" "TCGA-AZ-5407_334" "TCGA-AZ-6598_335"
#> [19] "TCGA-AZ-6599_336" "TCGA-AZ-6601_338" "TCGA-CA-6718_351"
#> [22] "TCGA-CI-6619_353" "TCGA-CI-6624_357" "TCGA-CK-4947_358"
#> [25] "TCGA-CK-4948_359" "TCGA-CK-4952_361" "TCGA-CK-5916_366"
#> [28] "TCGA-CL-5917_371" "TCGA-CM-4746_375" "TCGA-CM-5349_381"
#> [31] "TCGA-CM-5860_382" "TCGA-CM-5862_384" "TCGA-CM-6161_388"
#> [34] "TCGA-CM-6162_389" "TCGA-CM-6164_391" "TCGA-CM-6168_395"
#> [37] "TCGA-CM-6676_402" "TCGA-D5-6898_423" "TCGA-D5-6923_426"
#> [40] "TCGA-D5-6929_431" "TCGA-D5-6931_433" "TCGA-DC-5869_437"
#> [43] "TCGA-DC-6157_439" "TCGA-DC-6158_440" "TCGA-DC-6683_444"
#> [46] "TCGA-DM-A0XF_447" "TCGA-F4-6460_478" "TCGA-G4-6293_498"
#> [49] "TCGA-G4-6315_506" "TCGA-G4-6323_510" "TCGA-G4-6586_511"
#> [52] "TCGA-G5-6235_516"
```

The searching space can be also interactively explored as a collapsible tree or a sunburst.

```r
### visualising the CELLector searching space as a binary tree

CELLector.visualiseSearchingSpace(searchSpace = CSS,CLdata = CELLlineData)
```

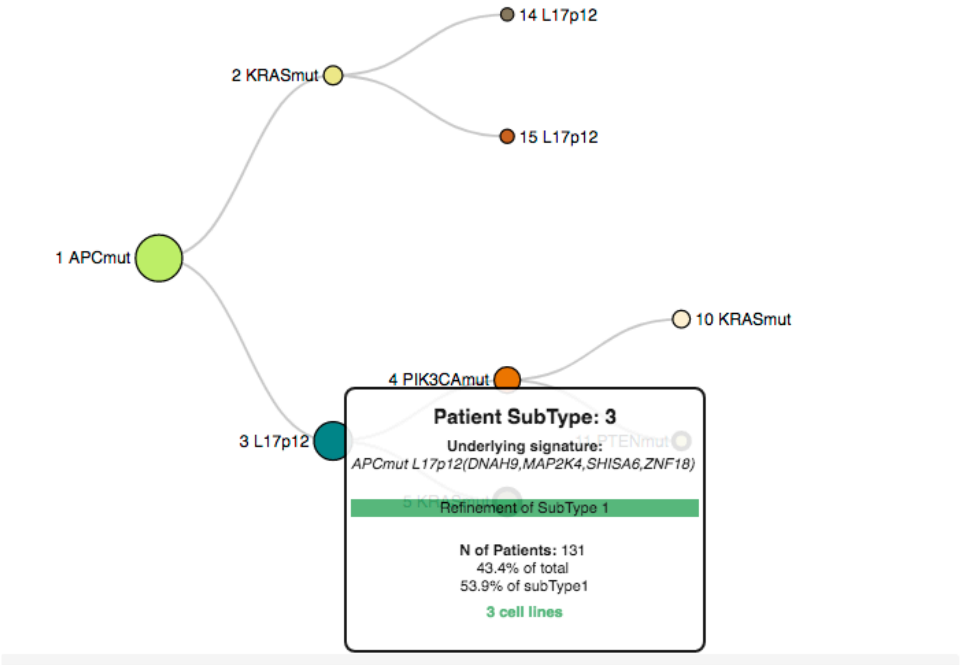

```
### visualising the CELLector searching space as a binary tree
CELLector.visualiseSearchingSpace_sunBurst(searchSpace = CSS)
```

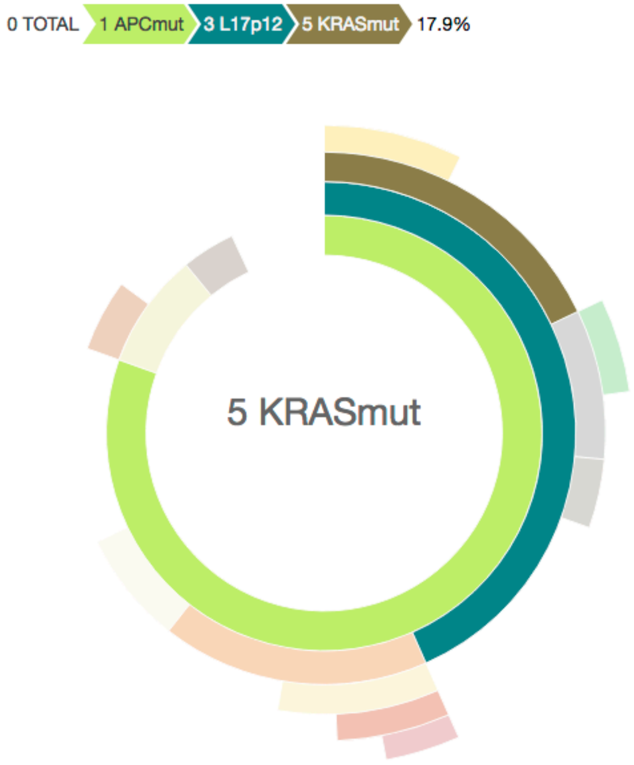

Finally 10 of the most representative cell lines can be selected with the following command:

```
### take all the signatures from the searching space
Signatures <- CELLector.createAllSignatures(CSS$navTable)
### mapping the cell lines on the CELLector searching space
ModelMat<-CELLector.buildModelMatrix(Signatures$ES,CELLlineData,CSS$navTable)

### selecting 10 cell lines
selectedCellLines<-CELLector.makeSelection(modelMat = ModelMat, n=10,
searchSpace = CSS$navTable)
selectedCellLines,align = 'l'
```

| Tumour.SubType.Index | Representative.Cell.Line | Signature | percentage.patients |
|---|---|---|---|
| 1 | CL-34 | APCmut | 80.463576 |
| 2 | SNU-C2B | ~APCmut KRASmut | 8.609272 |
| 3 | CCK-81 | APCmut L17p12(DNAH9,MAP2K4,SHISA6,ZNF18) | 43.377483 |
| 4 | CL-11 | APCmut ~L17p12(DNAH9,MAP2K4,SHISA6,ZNF18) PIK3CAmut | 17.218543 |
| 5 | GP5d | APCmut L17p12(DNAH9,MAP2K4,SHISA6,ZNF18) KRASmut | 17.880795 |
| 6 | LS-1034 | APCmut L17p12(DNAH9,MAP2K4,SHISA6,ZNF18) ~KRASmut PTENmut | 8.609272 |
| 7 | SW837 | APCmut L17p12(DNAH9,MAP2K4,SHISA6,ZNF18) KRASmut PIK3CAmut | 7.284768 |
| 8 | SNU-1040 | APCmut L17p12(DNAH9,MAP2K4,SHISA6,ZNF18) ~KRASmut ~PTENmut L5q21.1(…) | 3.973510 |
| 1 | COLO-205 | APCmut | 80.463576 |
| 2 | LS-180 | ~APCmut KRASmut | 8.609272 |

#### Scoring the clinical relevance of cancer cell lines

With CELLector is possible to quantify the quality of each cell line in terms of its ability to represent an entire cohort of considered disease matching patients. This is quantified as a trade-off between two factors. The first factor is the length of the CELLector signatures (in terms of number of composing individual alterations) that are present in the cell line under consideration. This is proportional to the granularity of the representative ability of the cell line, i.e. the longest the signature the more precisely defined is the represented sub-cohort of patients. The second factor is the size of the patient subpopulation represented by the signatures that can be observed in the cell line under consideration, thus accounting for the prevalence of the sub-cohort modeled by that cell line. An input parameter allows for these two factors to be weighted equally or differently.

In this example, we want to score the quality of the cell lines in the colorectal cancer panel, based on the searching space assembled in the previous examples.

```
### Scoring colorectal cancer cell lines based on the searching space assembled in the
previous examples

CSscores<-CELLector.Score(NavTab = CSS$navTable,CELLlineData = CELLlineData)

### Visualising the best 3 cell lines according to the considered searching space and
criteria
```

| CellLines | GlobalSupport | SignatureLength | CELLectorScores | Signature |
|---|---|---|---|---|
| SNU-1040 | 0.0927152 | 4 | 0.5231788 | APCmut, ~L17p12(DNAH9,MAP2K4,SHISA6,ZNF18), PIK3CAmut, PTENmut |
| COLO-678 | 0.0728477 | 4 | 0.5182119 | APCmut, ~L17p12(DNAH9,MAP2K4,SHISA6,ZNF18), ~PIK3CAmut, KRASmut |
| HCC2998 | 0.0728477 | 4 | 0.5182119 | APCmut, ~L17p12(DNAH9,MAP2K4,SHISA6,ZNF18), ~PIK3CAmut, KRASmut |

In this other example, we want to recompute the cell line scores but prioritising the size of the sub-cohort represented by each cell line over how much it is precisely defined.

```
### Scoring colorectal cancer cell lines based on the searching space assembled in the previous examples
CSscores<-CELLector.Score(NavTab = CSS$navTable,CELLlineData = CELLlineData,alfa = 0.10)

### Visualising the best 3 cell lines according to the considered searching space and criteria
```

| CellLines | GlobalSupport | SignatureLength | CELLectorScores | Signature |
|---|---|---|---|---|
| C2BBe1 | 0.8046358 | 1 | 0.7408389 | APCmut |
| CL-11 | 0.8046358 | 1 | 0.7408389 | APCmut |
| CL-34 | 0.8046358 | 1 | 0.7408389 | APCmut |

##### Assembling Genomic Binary Event Matrices (BEMs) and selecting cell line using customised genomic data

In the following example, we show how it is possible to run CELLector analyses and cell line selections from user defined genomic data. To this aim we will first generate genomic binary event matrices (BEMs) for Colorectal Cancer patients and cell lines using somatic variant catalogues derived from the TCGA (as presented in *Iorio et al, Cell 2016*) and from the latest installment of the *Cell Model Passports* (https://cellmodelpassports.sanger.ac.uk/), respectively for patients and cell lines, pre-selecting sets of cancer driver genes and variants to be considered in these catalogues. Fully user defined genomic variants catalogues can also be used. In this example we will particularly focus on **human derived microsatellite stable diploid colorectal carcinoma cell lines resected from male patients**.

First we build the BEMs as explained, with the following commands:

```
## loading a set of high-confidence cancer driver genes from Iorio et al, Cell 2016
data(CELLector.HCCancerDrivers)

## loading a set of variants observed it at least two patients in COSMIC
data(CELLector.RecfiltVariants)

## Assembling a genomic binary event matrix (BEM) for human derived microsatellite
stable diploid colorectal carcinoma cell lines,
## resected from male patients using genomic data from the Cell Model Passports,
considering only variants observed in COSMIC in at lest two patients
## in high-confidence cancer driver genes
COREAD_cl_BEM <- CELLector.CELLline_buildBEM(
                  Tissue='Large Intestine',
                  Cancer_Type = 'Colorectal Carcinoma',
                  msi_status_select = 'MSI',
                  ploidy_th = c(2,2),
                  GenesToConsider = CELLector.HCCancerDrivers,
                  VariantsToConsider = CELLector.RecfiltVariants)
```

```
## bar diagram with mutation frequencies in the BEM for 20 top frequently mutated genes
barplot(sort(colSums(COREAD_cl_BEM[,3:ncol(COREAD_cl_BEM)]),decreasing=TRUE)[1:20],
        las=2,ylab='n. mutated cell lines')
```

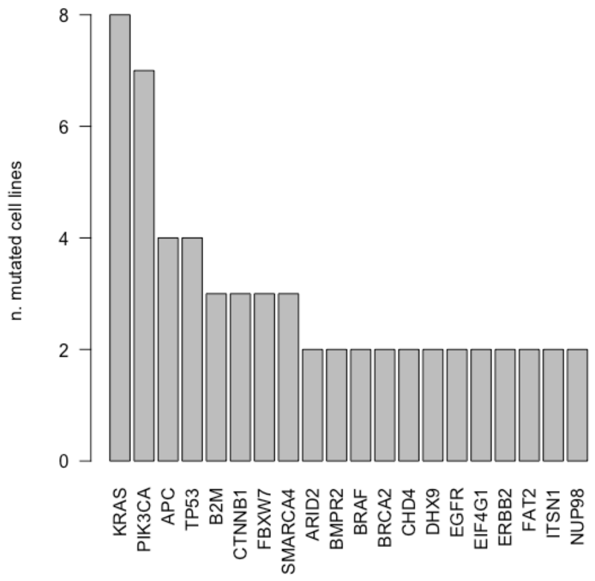

```
## Assembling BEM for Colorectal Adenocarcinoma (COAD/READ) primary tumours using data from the TCGA as presented in Iorio et al 2016,
## considering only variants observed in COSMIC in at lest two patients in high-confidence cancer driver genes
COREAD_tum_BEM<-
  CELLector.Tumours_buildBEM(Cancer_Type = 'COAD/READ',GenesToConsider = CELLector.HCCancerDrivers,
                             VariantsToConsider = CELLector.RecfiltVariants)

## showing a bar diagram with mutation frequency of 30 top frequently altered genes
barplot(100*sort(rowSums(COREAD_tum_BEM),
                 decreasing=TRUE)[1:30]/ncol(COREAD_tum_BEM),
                 las=2,ylab='% patients')
```

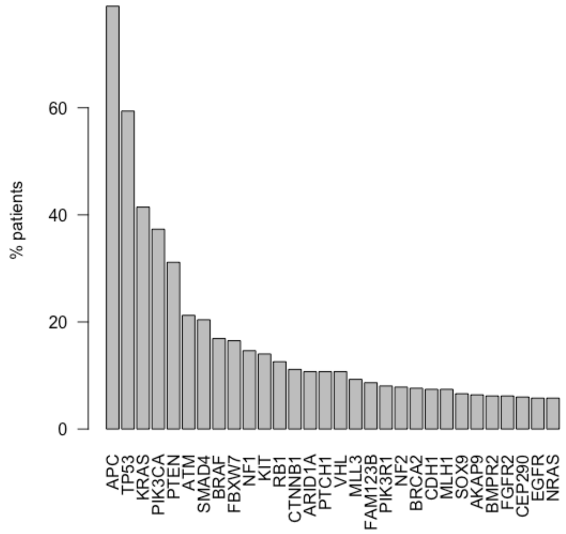

Next, we assemble a CELLector searching space and select representative cell lines as for the previous examples.

```
### Assembling the searching space and visualising it as interactive sunburst

COREAD_tum_BEM<-CELLector.unicizeSamples(COREAD_tum_BEM)
CSS<-CELLector.Build_Search_Space(ctumours = t(COREAD_tum_BEM),
                                  verbose = FALSE,
                                  minGlobSupp = 0.02,
                                  cancerType = 'COREAD')
### take all the signatures from the searching space
Signatures <- CELLector.createAllSignatures(CSS$navTable)

### mapping the cell lines on the CELLector searching space
ModelMat<-CELLector.buildModelMatrix(Signatures$ES,COREAD_cl_BEM,CSS$navTable)

### selecting 5 representative cell lines
selectedCellLines<-CELLector.makeSelection(modelMat = ModelMat,
                                           n=5,
                                           searchSpace = CSS$navTable)
```

| Tumour.SubType.Index | Representative.Cell.Line | Signature | percentage.patients |
|---|---|---|---|
| 1 | LoVo | APCmut | 78.96907 |
| 2 | KM12 | ~APCmut, TP53mut | 10.92784 |
| 3 | RKO | APCmut, TP53mut | 48.45361 |
| 4 | HCT-15 | APCmut, ~TP53mut, KRASmut | 14.02062 |
| 5 | SNU-C2B | APCmut, TP53mut, KRASmut | 20.20619 |

### 4. CELLector R Shiny App Tutorial

The CELLector app can be accessed at https://ot-cellector.shinyapps.io/cellector_app/. The user interface of the CELLector app has two tabs: respectively labelled with *Select Cell Lines* and *BEM builder*.

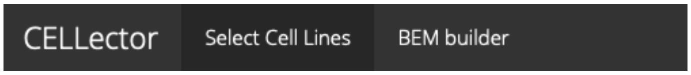

To run CELLector analyses and explore the results the first tab *Select Cell Lines* tab should be clicked (it is by default). By clicking on the second tab (*BEM builder*) a module for the generation of customised genomic binary event matrices (BEMs) for cell lines and tumours can be accessed (details in the sections below).

#### 1. ‘*Select Cell Lines*’ TAB

The ‘*Select Cell Lines*’ page displays seven boxes, detailed in the sections below.

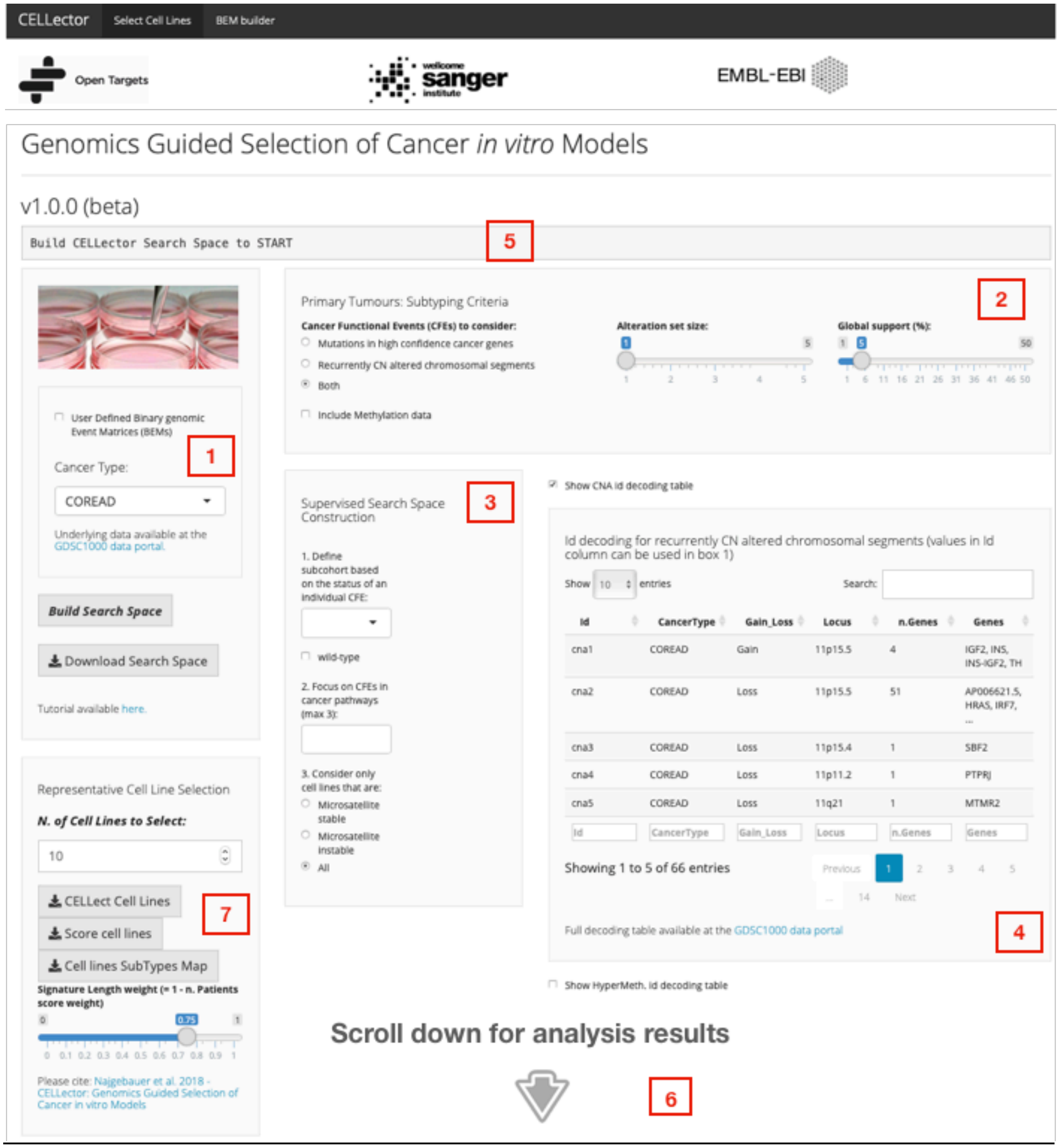

#### Step-by-step User Guide

In this tutorial, we demonstrate how to set up and explore the CELLector analysis. For this purpose, we reproduce the example described in the Case Study 1.

#### Case Study 1

Briefly, we want to select 5 most clinically relevant *in vitro* models that best represent the genomic diversity of *TP53* mutated colorectal tumours. The models we wish to select should be microsatellite stable and harbour at least one alteration in the following signalling pathways: PI3K-AKT-MTOR signalling, RAS-RAF-MEK-ERK/JNK signalling and WNT signalling. Finally, we want the model selection to be guided based on somatic mutations and copy number alterations that are prevalent in at least 3% of the studied *TP53* mutant tumour cohort. The summary results from this analysis are shown in Case Study Figure 1.

**A.**

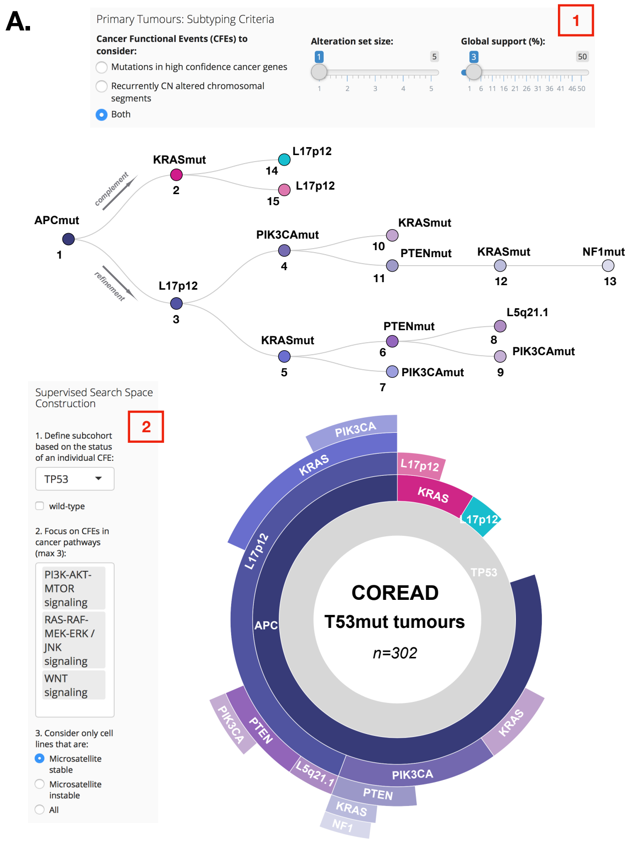

**B.**

~grey - absence
**bold** - presence

***SELECT n=5***

| Sub Type | Genomic Signature | n. Patients | % | Representative Cell Lines |
|---|---|---|---|---|
| 1 | **APCmut** | 243 | 80.46 | C2BBe1, CL-11, CL-34, CL-40, **COLO-320-HSR**, DIFI, HCC2998, HT-115, HT-29, HT55, LS-1034, LS-123, NCI-H747, RCM-1, SNU-1040, SNU-61, SNU-81, SW1116, SW1417, SW1463, SW620, SW837, SW948, T84 |
| 2 | ~APCmut **KRASmut** | 26 | 8.61 | **HCC-56** |
| 14 | ~APCmut ~KRASmut **L17p12(DNAH9,MAP2K4,SHISA6,ZNF18)** | 12 | 3.97 | ***lack of MSS in vitro models*** |
| 3 | **APCmut L17p12(DNAH9,MAP2K4,SHISA6,ZNF18)** | 131 | 43.38 | CL-11, SW620, **SW837** |
| 15 | ~APCmut **KRASmut L17p12(DNAH9,MAP2K4,SHISA6,ZNF18)** | 14 | 4.64 | ***lack of MSS in vitro models*** |
| 4 | **APCmut** ~L17p12(DNAH9,MAP2K4,SHISA6,ZNF18) **PIK3CAmut** | 52 | 17.22 | CL-40, **HT-115**, SNU-1040, SW948, T84 |
| 10 | **APCmut** ~L17p12(DNAH9,MAP2K4,SHISA6,ZNF18) ~PIK3CAmut **KRASmut** | 22 | 7.28 | HCC2998, LS-1034, LS-123, NCI-H747, RCM-1, SNU-61, **SNU-81**, SW1116, SW1463 |
| 5 | **APCmut L17p12(DNAH9,MAP2K4,SHISA6,ZNF18) KRASmut** | 54 | 17.88 | CL-11, SW620, SW837 |
| 11 | **APCmut** ~L17p12(DNAH9,MAP2K4,SHISA6,ZNF18) **PIK3CAmut PTENmut** | 28 | 9.27 | SNU-1040 |
| 7 | **APCmut L17p12(DNAH9,MAP2K4,SHISA6,ZNF18) KRASmut PIK3CAmut** | 22 | 7.28 | ***lack of MSS in vitro models*** |
| 12 | **APCmut** ~L17p12(DNAH9,MAP2K4,SHISA6,ZNF18) **PIK3CAmut PTENmut KRASmut** | 18 | 5.96 | ***lack of MSS in vitro models*** |
| 6 | **APCmut L17p12(DNAH9,MAP2K4,SHISA6,ZNF18)** ~KRASmut **PTENmut** | 26 | 8.61 | ***lack of MSS in vitro models*** |
| 8 | **APCmut L17p12(DNAH9,MAP2K4,SHISA6,ZNF18)** ~KRASmut ~PTENmut **L5q21.1(APC...)** | 12 | 3.97 | ***lack of MSS in vitro models*** |
| 13 | **APCmut** ~L17p12(DNAH9,MAP2K4,SHISA6,ZNF18) **PIK3CAmut PTENmut KRASmut NF1mut** | 11 | 3.64 | ***lack of MSS in vitro models*** |
| 9 | **APCmut L17p12(DNAH9,MAP2K4,SHISA6,ZNF18)** ~KRASmut **PTENmut PIK3CAmut** | 15 | 4.97 | ***lack of MSS in vitro models*** |

**Case Study Figure 1 CELLector analysis results: Case Study 1.**
**A.** Visual representation of the CELLector search space summarising the genomic heterogeneity of *TP53* mutant colorectal (COREAD) tumours, considering both somatic mutations and copy number alterations in the user-defined biological pathways (user interface box 1 and box 2). Each node of the binary tree (top) represents a tumour subpopulation with define genomic signature.

The prevalence of the identified signatures, and their hierarchical co-occurrence is represented by the sunburst (below). Each segment of the sunburst corresponds to a node in the tree and is color-coded accordingly. The CELLector search space is built using genomic characterisation of a cohort of 517 colorectal cancer patient available as a built-in dataset (Table S1). First, the cohort is reduced to the 302 tumours harbouring *TP53* mutations. CELLector then identifies 3 major molecular subpopulations characterised, respectively, by *APC* mutations, *KRAS* mutations, and loss of a segment of chromosome 17 (at the 17p12 loci) that collectively represent 93 % (n=243+26+12 = 281) of the studied tumour cohort. The remaining 7% of the *TP53* mutant tumours (n=21) do not fall into any of the identified molecular subpopulations thus are not represented in the CELLector search space. The largest molecular subpopulation (80.46%, n=243, harbouring *TP53* and *APC* mutations) is assigned to the root of the search space (node 1, in purple). The second largest subpopulation (9%, n=26) is characterised by co-occurrence of *TP53* and *KRAS* mutations in absence of *APC* mutations (node 2, in magenta), and the third largest subpopulation (4%, n=12) harbours *TP53* mutations and the loss of 17p12 segment in the absence of both *APC* and *KRAS* mutations (node 14, in cyan). Each of these complementary tumour subpopulations are then further refined based on the prevalence of the remaining set of molecular alterations (as described in STAR Methods). This process runs recursively and stops when all molecular alteration sets with user-determined tumour prevalence (in this case 3%, user interface box 1) are defined. In this study case, a total number of 15 distinct tumour subpopulations with defined genomic signatures are identified, when considering the *TP53* mutant tumour sub-cohort and alterations in the selected pathways. **B.** Cell Line Map table including microsatellite stable cell lines mirroring the genomic signatures of the *TP53* mutant COREAD tumour subpopulations identified in the CELLector search space (user interface box 2). The Cell Map table uncovers the complete set of molecular alterations (e.g. genomic signatures) that characterise each tumour subpopulation. For example, the least prevalent *TP53* mutant colorectal cancer subpopulation (node 13, 3.64% of patients) is characterised by co-occurring mutations in *APC, PIK3CA, PTEN, KRAS* and *NF1* in the absence of 17p12 segment loss; this genomic signature is not mirrored by any of the considered microsatellite stable (MSS) colorectal cancer models. The representative cell lines are picked from each of the molecular tumour subpopulations starting with the most prevalent one (as detailed in the STAR Methods). The models in green represent a possible choice of *n*-user-defined cell lines that could be selected in the presented case study.

#### Step 1: Setting Up the Searching space building criteria

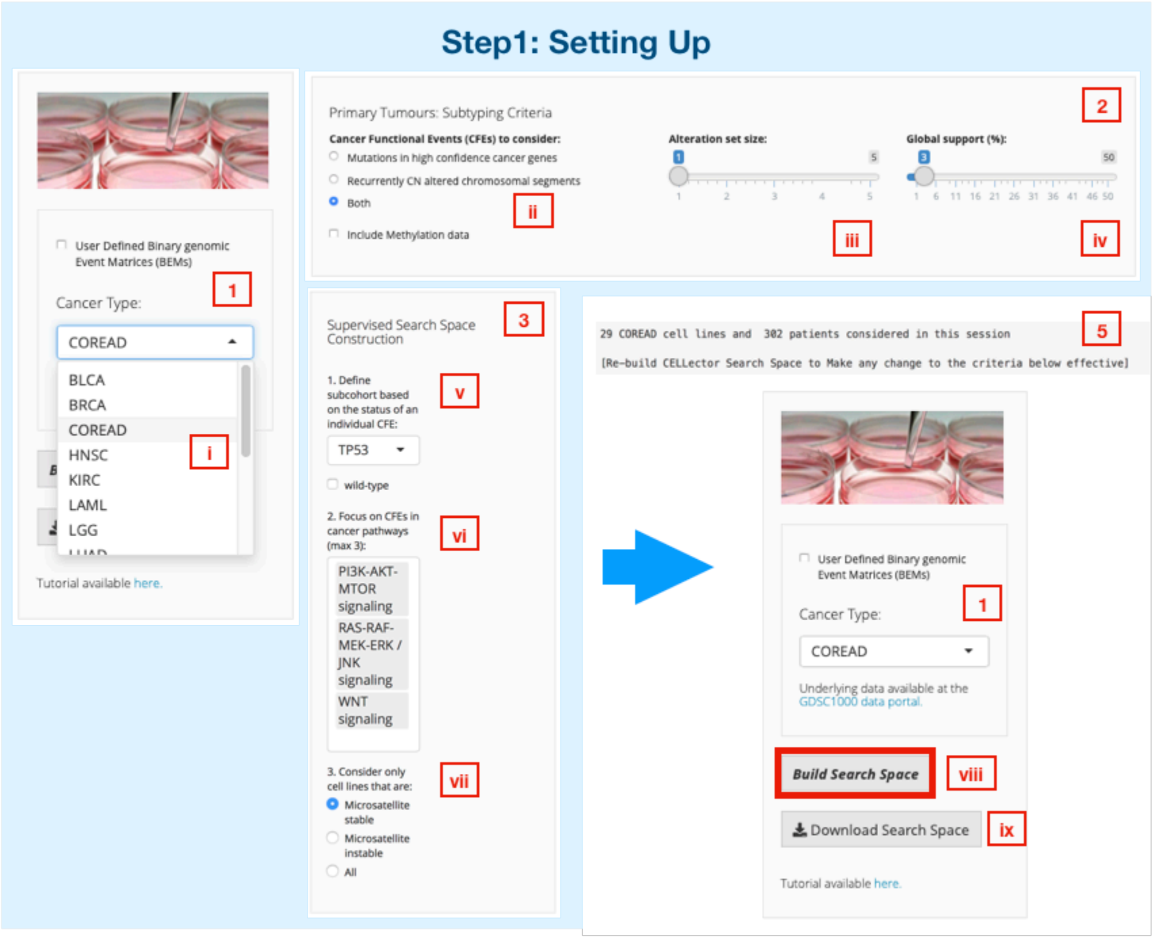

##### *Box 1: Cancer Type*

CELLector provides built-in genomics data for disease-matched primary tumours and cell lines derived from 16 cancer types (Table S1). To start, select a cancer type of interest from a dropdown list (as shown above) or alternatively, upload your own reference data by ticking the '*User Defined Binary genomic Event Matrices (BEMs)*' box (as shown below). In this example case study, we wish to select colorectal models (COREAD) (i).

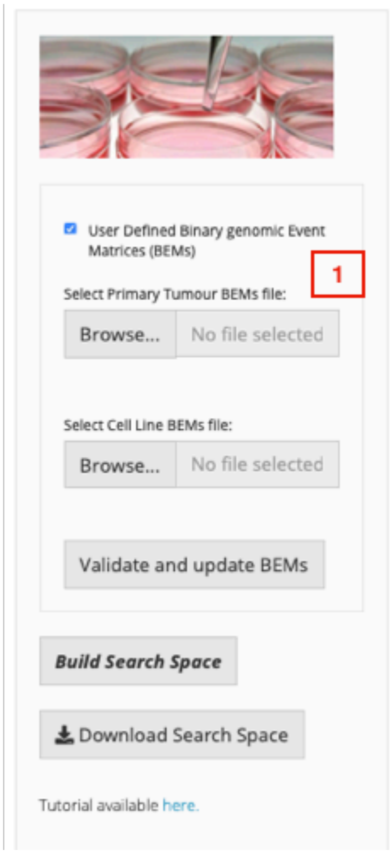

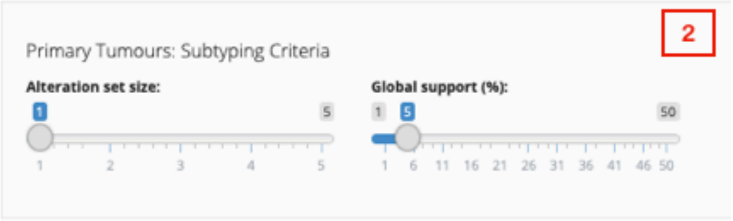

##### *Box 2: Primary Tumours: Subtyping Criteria*

Before running the analysis, user has to specify some parameters that determine the way the CELLector search space is constructed. Otherwise the analysis runs with the default settings. In this example case study, we want to represent genomic diversity of the studied cohort based on both somatic mutations and copy number alterations (ii). Alteration set size is set to 1 (default setting; iii), meaning that the algorithm looks for only one alteration at the time that is the most prevalent in recursively identified tumour subpopulations. If the alteration set size were set up to 2 (max 5), the algorithm would look for a co-occurring combination of two (up to 5) alterations at the time in recursively identified tumour subpopulations. Lastly, the genomic signatures (e.g. the combination of co-occurring genomic alterations) underlying the identified tumour subpopulations have to be prevalent in at least 3% of the studied tumour cohort (iv).

These steps (ii, iii and iv) are essential to build the CELLector search space and subsequently map representative *in vitro* models. If no other criteria are specified (see below, Box 3: Supervised Search Space Construction), the analysis runs on the whole cohort of COREAD tumours (n=517) and entire set of COREAD cell lines (n=51) available *via* CELLector (Table S1).

When user defined reference data (Box 1) is used, the following parameters: *alteration set size* and *global support* (Box 2) have to be specified before running the analysis (as shown above).

##### *Box 3: Supervised Search Space Construction*

To flexibly tailor the model selection to fit the context of a study, users can restrict the tumour cohort based on the status of an individual alteration (v), focus on alterations in cancer pathways (vi) or/and consider microsatellite status of cell lines (vii).

In this case study, we want to select cell lines that best represent a colorectal cancer patient sub-cohort that harbours *TP53* mutation. In addition, models should harbour at least one alteration in the following signalling pathways: PI3K-AKT-MTOR signalling, RAS-RAF-MEK_ERK/JNK signalling and WNT signalling and be microsatellite stable.

##### *Box 1: Build Search Space*

Once the desire criteria (Box 2 and Box 3 if build-in datasets are used or Box 2 if user defined reference data are used) are defined, we can start the analysis by clicking '*Build Search Space*' button (viii). The information regarding the number of primary tumours and cell lines considered in the CELLector analysis is displayed in Box 5. The CELLector search space and the cell line selection results are shown in Box 6 (see below Step 2: Exploring the CELLector Search Space). The CELLector Search Space is translated into a Cell Line Map table (as shown in Case Study Figure 1B) and can be downloaded as text file to the user specified location (ix).

##### *Box 4: cna and hms id decoding*

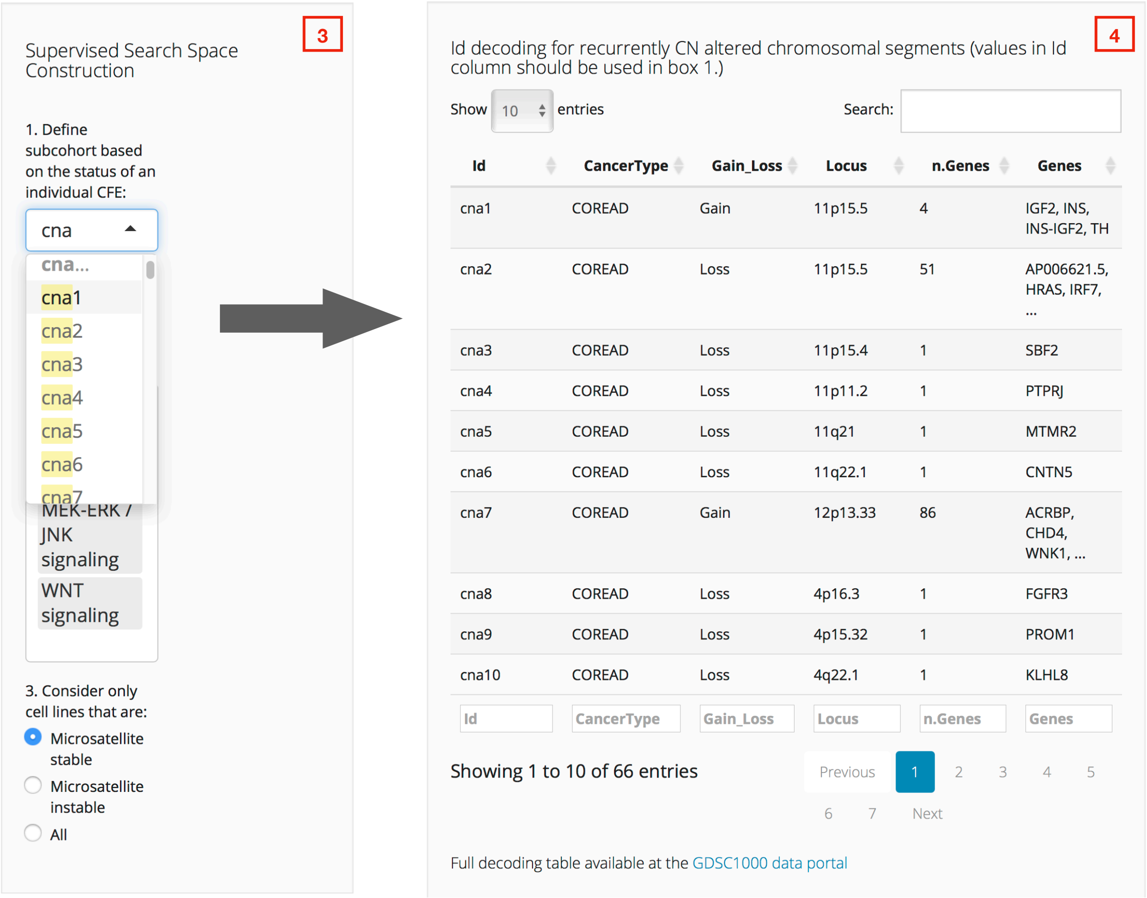

If a user wishes to include copy number alterations (can) or hypermethylation (hsm) in the analysis (Box 3: Supervised Search Space Construction), CELLector provides a look up tables to search for genes that are within recurrently copy number altered chromosomal segments (Box 4) or hypermethylated regions (table not shown).

#### Step 2: Exploring the CELLector Search Space

CELLector supports a range of visualisation tools that enable users to interactively explore the CELLector search space (e.g. the identified tumour subpopulations with defined genomic signature) and the cell line selection results (Box 6). These visualisation tools include binary tree (A), sunburst (B), pie chart (C), dot plots (D) and cell line information table (E).

6

#### Step2: Exploring the CELLector Search Space

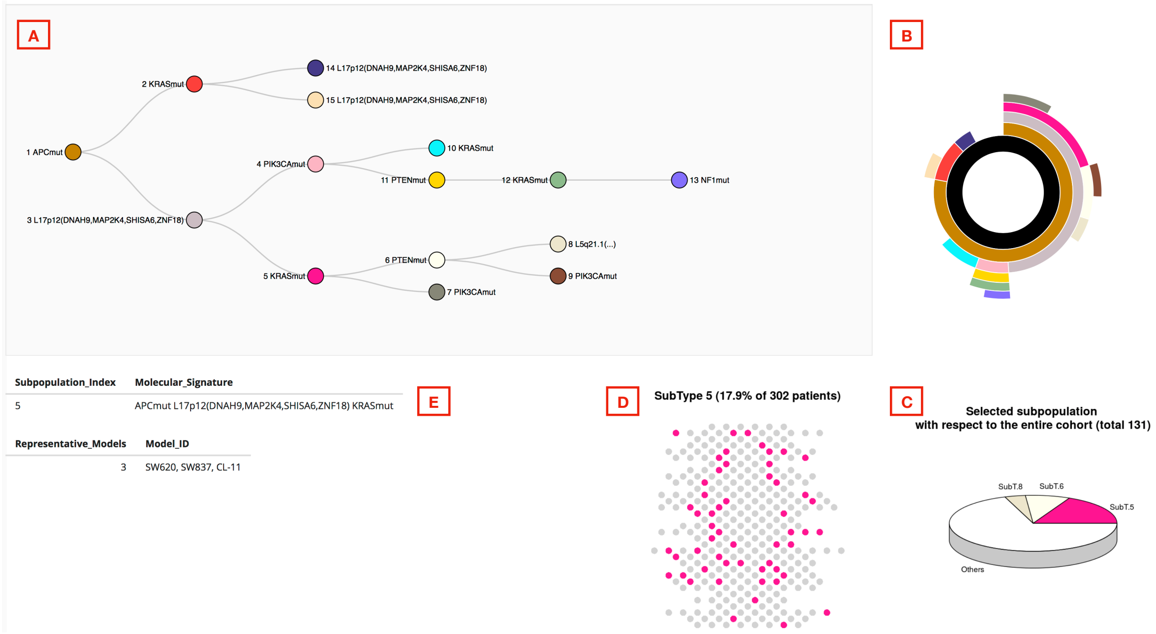

A binary tree, which represents the genomic diversity of the examined patient cohort, pops up in the bottom of the page (after clicking on ‘*Build Search Space*’ button, box 1). Starting from the root (the largest tumour subpopulation), two sibling nodes are added. Upper node represents complementary subpopulation, while lower node represents refinement of the parent subpopulation (root). To support an easy interpretation of the resulting binary tree, the considered cohort is represented as sunburst that takes into account the prevalence of the identified genomic signatures and their hierarchical co-occurrence (B). Users can interactively explore this tree by clicking on its nodes (A). Nodes with a bold circle are expandable (not shown). After clicking on a node, the tumour subpopulation information relating to this node is then displayed (information relating to node 5, in magenta, is shown). The segments of the sunburst (B), dot plot (D) and pie chart (C), which correspond to selected tumour subpopulation, are color-coded accordingly. The molecular signature of a tumour subpopulation is shown below the tree, together with the information regarding the number and names of cell line models that mirror this signature (E). This example subpopulation (node 5, in magenta) is defined by the co-occurring combination of mutations in *TP53, APC*, *KRAS* and rearrangement in segment *L17p12* and covers 17.9% of *TP53* mutated tumours (as shown in E and D).

There are 3 cell line models that mirror this molecular signature (*SW620, SW837, CL-11*, as shown in E).

To enable easy navigation, the resulting CELLector searching space is translated into a Cell Map table (as shown in Case Study Figure 1B), which can be downloaded by clicking ‘Download Search Space’ (Box 1, iv).

##### Step 3: Model Selection

The last step in the CELLector analysis, after constricting search space (Step 1), is the selection of the user defined number of representative cell line models (Box 7).

The list of possible model choices (in this case list of 5 representative models) is downloaded to the user-specified directory by clicking ‘*CELLect Cell Lines*’. The file provides information regarding the prevalence of the tumour subpopulations the selected model(s) represent together with corresponding genomic signatures. The ‘*Cell lines SubTypes Map*’ can be also downloaded to the user-specified directory by clicking the ‘*Cell lines SubTypes Map*’. Finally, the user can score the cell lines by specifying the weight of signature length (bottom of Box 7). The list with cell line scores is downloaded by clicking ‘*Score cell lines*’.

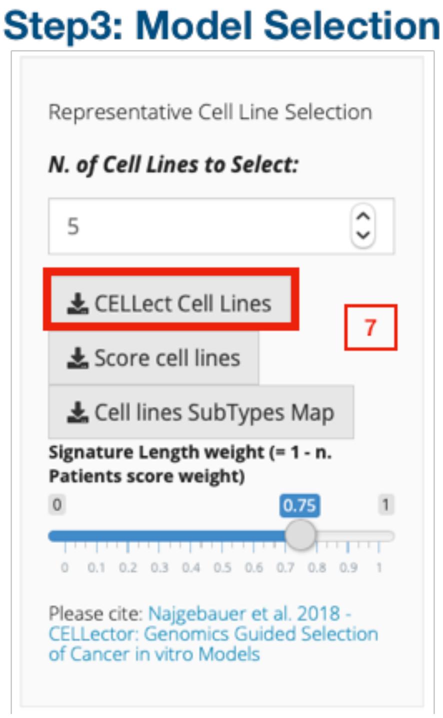

#### Case Study 2

In this case study, we want to select 5 microsatellite stable cell lines capturing the genomic heterogeneity of a large cohort of colorectal cancer patients, focusing on sets of somatic mutations that are prevalent in at least 5% of the considered patient cohort. The summary results from this analysis are shown in Case Study Figure 2.

A.

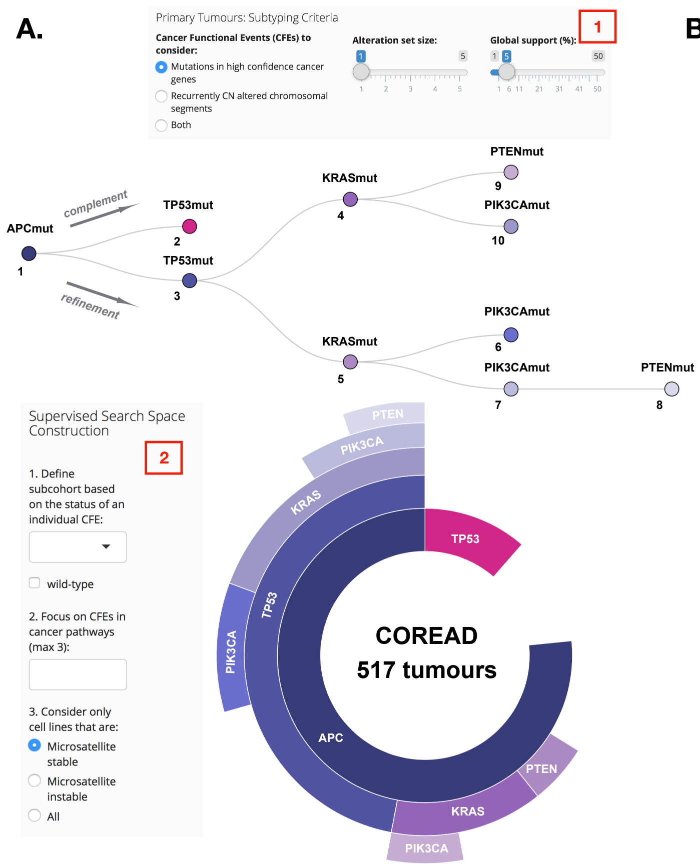

B.

~grey - absence
**bold** - presence

***SELECT n=5***

| Sub Type | Genomic Signature | n Patients | % | Representative Cell Lines |
|---|---|---|---|---|
| 1 | **APCmut** | 396 | 76.60 | C2BBe1, CL-11, CL-34, CL-40, COLO-205, COLO-320-HSR, COLO-678, DIFI, HCC2998, HT-115, HT-29, HT55, LS-1034, LS-123, **MDST8**, NCI-H747, RCM-1, SK-CO-1, SNU-1040, SNU-61, SNU-81, SW1116, SW1417, SW1463, SW620, SW837, SW948, T84 |
| 2 | ~APCmut **TP53mut** | 59 | 11.41 | CaR-1, HCC-56, NCI-H508, **NCI-H716**, SNU-C1 |
| 3 | **APCmut TP53mut** | 243 | 47.00 | C2BBe1, CL-11, CL-34, CL-40, COLO-320-HSR, DIFI, HCC2998, HT-115, HT-29, **HT55**, LS-1034, LS-123, NCI-H747, RCM-1, SNU-1040, SNU-61, SNU-81, SW1116, SW1417, SW1463, SW620, SW837, SW948, T84 |
| 4 | **APCmut** ~TP53mut **KRASmut** | 71 | 13.73 | COLO-678, **SK-CO-1** |
| 9 | **APCmut** ~TP53mut ~KRASmut **PTENmut** | 28 | 5.42 | ***lack of microsatellite stable in vitro models*** |
| 5 | **APCmut TP53mut KRASmut** | 100 | 19.34 | CL-11, CL-40, HCC2998, **LS-1034**, LS-123, NCI-H747, RCM-1, SNU-61, SNU-81, SW1116, SW1463, SW620, SW837, SW948, T84 |
| 10 | **APCmut** ~TP53mut **KRASmut PIK3CAmut** | 31 | 6.00 | ***lack of microsatellite stable in vitro models*** |
| 6 | **APCmut TP53mut** ~KRASmut **PIK3CAmut** | 52 | 10.06 | HT-115, SNU-1040 |
| 7 | **APCmut TP53mut KRASmut PIK3CAmut** | 46 | 8.90 | CL-40, SW948, T84 |
| 8 | **APCmut TP53mut KRASmut PIK3CAmut PTENmut** | 27 | 5.22 | ***lack of microsatellite stable in vitro models*** |

#### Case Study Figure 2 CELLector analysis results: Case Study 2.

**A.** Visual representation of the CELLector search space constructed based on the prevalence of co-occurring mutations in a cohort of colorectal cancer (COREAD) patients (user interface box 1). Each node of the binary tree (top) represents a tumour subpopulation with a defined genomic signature. The prevalence of the identified signatures, and their hierarchical co-occurrence is represented by the sunburst (below). Each segment of the sunburst corresponds to a node in the three and is color-coded accordingly. The CELLector search space is assembled using a built-in dataset containing the genomic characterisation of a cohort of 517 colorectal cancer tumours (Table S1). CELLector identifies 2 major subpopulations defined by *APC* mutations (node 1, in purple) and *TP53* mutations (node 2, in magenta), collectively representing 88% of the studied cohort (n = 396 + 59 = 455 patients). The remaining 12% of tumours (n = 21) do not fall into any of the identified molecular subpopulations, i.e. they do not harbour *APC* nor *TP53* mutations. The largest molecular subpopulation (76.6%, n = 396, harbouring *APC* mutations) is assigned to the root of the search space (node 1). The second largest subpopulation (11.4%, n=59) is characterised by *TP53* mutations in the absence of *APC* mutations (node 2). At this point, each identified tumour subpopulation is further refined based on the prevalence of other set of alterations (STAR Methods). This process runs recursively and stops when all alteration sets with a user-determined prevalence (in this case 5%, user interface box 1) have been identified. In this case study, a total number of 10 distinct tumour subpopulations with corresponding genomic signatures are identified. Notably, some of the identified signatures (such as absence of *APC* mutations (~APCmut), co-occurrence of mutations in *APC* and *TP53* (APCmut TP53mut), same combination with the addition of *KRAS* mutations (APCmut TP53mut KRASmut), and the co-occurrence of *APC* and *KRAS* mutations (APCmut ~TP53mut KRASmut) in the absence of *TP53* mutations were reported to have a prognostic role in colorectal tumour stratification (Schell et al., 2016). **B.** Cell Line Map table including microsatellite stable cell lines mirroring the genomic signatures of the COREAD subpopulations identified in the CELLector search space (user interface box 2). The Cell Map table uncovers the complete set of molecular alterations (e.g. genomic signatures) defining each tumour subpopulation. For example, the least prevalent colorectal cancer subpopulation (node 8, 5.22% of tumours) is characterised by the co-occurrence of *APC, TP53, KRAS, PIK3CA* and *PTEN* mutations; this genomic signature is not reflected by any of the available microsatellite stable COREAD models included in the built-in dataset. The representative cell lines are picked from each of the molecular tumour subpopulations (STAR Methods) starting from the most prevalent one. The models in green represent a possible choice of *n*-user-defined cell lines that could be selected in the presented case study.

#### 2. *'BEM builder' TAB*

The '*BEM Builder*' page displays '*Primary tumours*' and '*in-vitro models*' panels (as shown below), which are described in the relevant sections below.

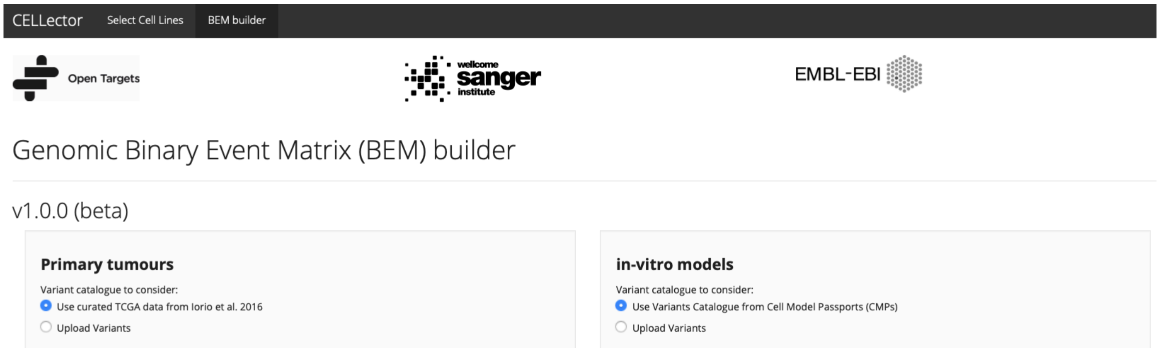

#### Primary tumours BEM builder

The primary tumours BEM can be generated using variant catalogue from either curated TCGA data (Iorio et al. 2016) or user own reference data.

##### Build-in TCGA data

To generate tumour BEM from build-in TCGA data select the '*use curated TCGA data from Iorio et al. 2016*' (step 1) then select cancer type of interest (step 2) and determine what genes and variants to consider (step 3). Supported options are *'All'* genes/variants, *'Iorio et al.2016'* drivers/variants, Cell Model Passports *'CMP drivers'* and *'User defined list'* for both genes and variants. Once all the parameters are specified, select BEM file format (R object or .tsv) and click '*Make new BEM*' (step 4). The information regarding the number of tumours and mutated genes used to generate the BEM will display in a box below (as shown below). Finally, once the BEM is generated, it is saved to user specified directory by clicking '*Save BEM*' (step 5).

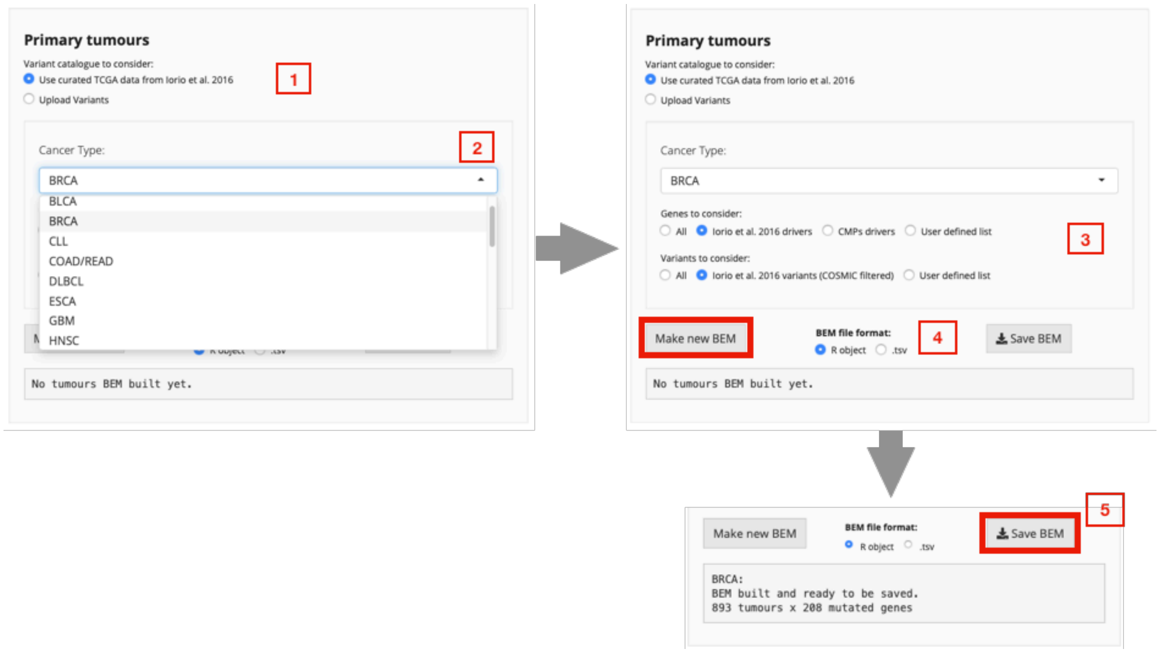

##### User defined tumour data

To generate tumour BEM using user defined tumour data select the '*Upload Variants*' (step 1) then select the data from your local directory (step 2), select BEM file format (R object or .tsv) and click '*Make new BEM*' (step 3). The information regarding the number of tumours and mutated genes used to generate the BEM will display in a box below. Finally, once the BEM is generated, it is saved to user specified directory by clicking '*Save BEM*' (step 4).

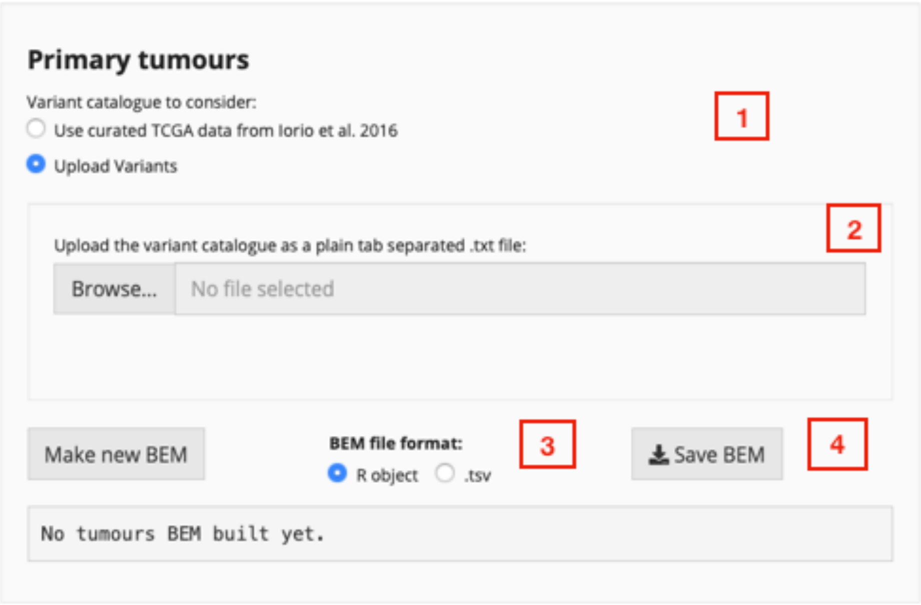

#### *In vitro* models BEM builder

The *in vitro* models BEM can be generated using variant catalogue from either Cell Model Passports (CMPs) or user own reference data.

##### Cell Model Passports data

To generate *in vitro* models BEM using variant catalogue from Cell Model Passports (CMPs) select the '*use Variant Catalogue from Cell Model Passports (CMPs)*' (step 1) then determine what types of models you are interested in (step 2). Once all the parameters are specified, select tissue of interest (step 3). The information about available models for that tissue type based on applied filters is display on the top of the panel (step 3).

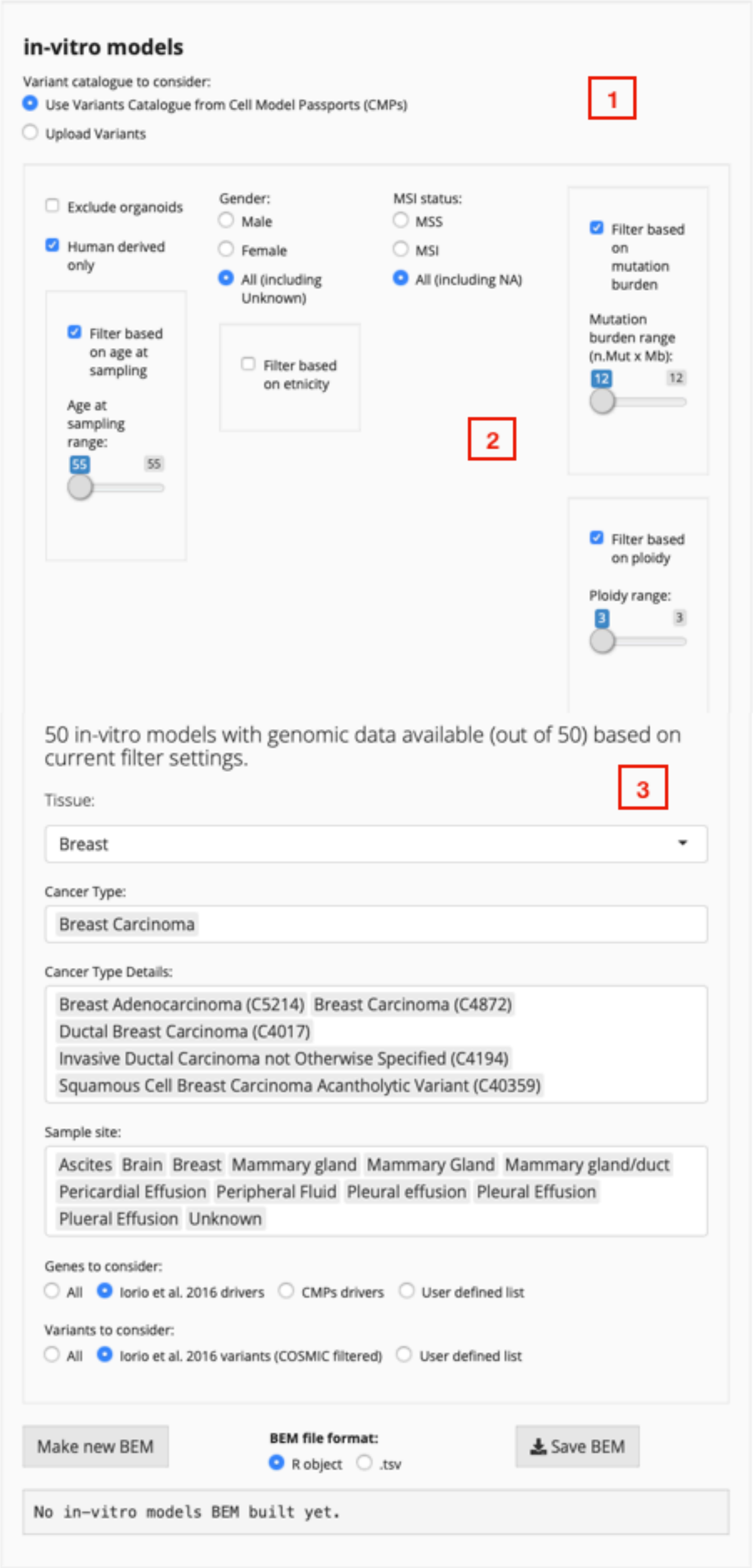
in-vitro models
Variant catalogue to consider:
Use Variants Catalogue from Cell Model Passports (CMPs)
Upload Variants
1
Exclude organoids
Human derived only
Filter based on age at sampling
Age at sampling range:
55
55
Gender:
Male
Female
All (including Unknown)
Filter based on etnicity
MSI status:
MSS
MSI
All (including NA)
2
Filter based on mutation burden
Mutation burden range (n.Mut x Mb):
12
12
Filter based on ploidy
Ploidy range:
3
3
50 in-vitro models with genomic data available (out of 50) based on current filter settings.
3
Tissue:
Breast
Cancer Type:
Breast Carcinoma
Cancer Type Details:
Breast Adenocarcinoma (C5214)
Breast Carcinoma (C4872)
Ductal Breast Carcinoma (C4017)
Invasive Ductal Carcinoma not Otherwise Specified (C4194)
Squamous Cell Breast Carcinoma Acantholytic Variant (C40359)
Sample site:
Ascites
Brain
Breast
Mammary gland
Mammary Gland
Mammary gland/duct
Pericardial Effusion
Peripheral Fluid
Pleural effusion
Pleural Effusion
Plueral Effusion
Unknown
Genes to consider:
All
Iorio et al. 2016 drivers
CMPs drivers
User defined list
Variants to consider:
All
Iorio et al. 2016 variants (COSMIC filtered)
User defined list
Make new BEM
BEM file format:
R object
.tsv
Save BEM
No in-vitro models BEM built yet.

Within the selected tissue, cancer type and cancer type details can be further defined. In the example below, we demonstrate how to generate models BEM for *breast carcinoma* cell lines derived from *ascites, breast, mammary gland, peripheral fluid and pleural effusion*. Applying these criteria, the number of available models was reduced from 50 (shown above, top panel step 3) to 10 (shown below, top panel step 3). Then we need to specify what genes and variants to consider. Supported options are *'All'* genes/variants*, 'Iorio et al.2016'* drivers/variants*,* Cell Model Passports *'CMP drivers'* and *'User defined list'* for both genes and variants. Once all the parameters are specified, select BEM file format (R object or .tsv) and click '*Make new BEM*' (step 4). The information regarding the number of tumours and mutated genes used to generate the BEM will display in a box below (as shown below). Finally, once the BEM is generated, it is saved to user specified directory by clicking '*Save BEM*' (step 5).

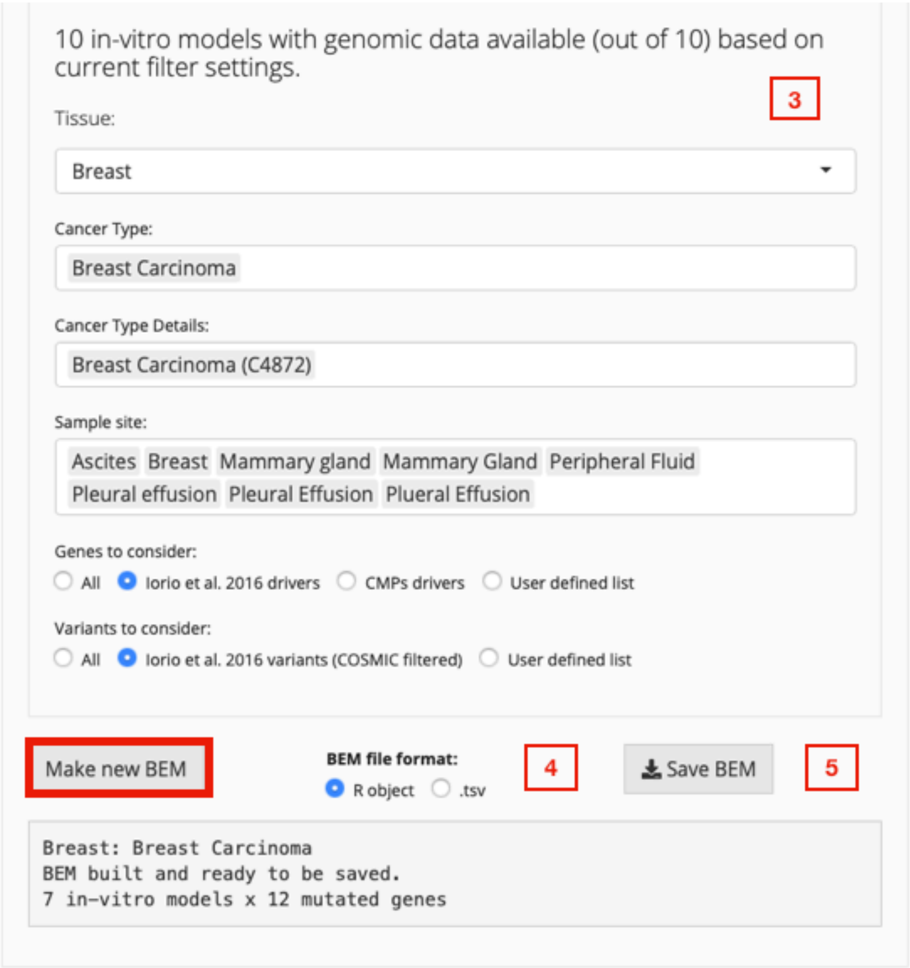

##### User defined model data

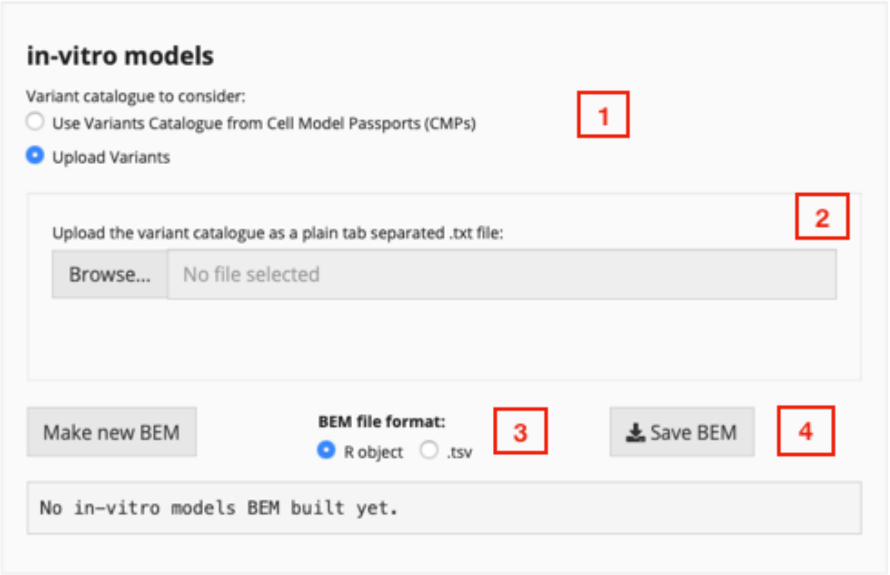

To generate *in vitro* models BEM using user defined model data select the '*Upload Variants*' (step 1) then select the data from your local directory (step 2), select BEM file format (R object or .tsv) and click '*Make new BEM*' (step 3). The information regarding the number of models and mutated genes used to generate the BEM will display in a box below. Finally, once the BEM is generated, it is saved to user specified directory by clicking '*Save BEM*' (step 4).

### 5. Running the CELLector R Shiny App Locally

Download the CELLector Rshiny App code as a single compressed folder from https://github.com/francescojm/CELLector_App/archive/master.zip.
Uncompress the folder, open Rstudio (available here https://www.rstudio.com/) and set the working directory to that folder (see below figure).

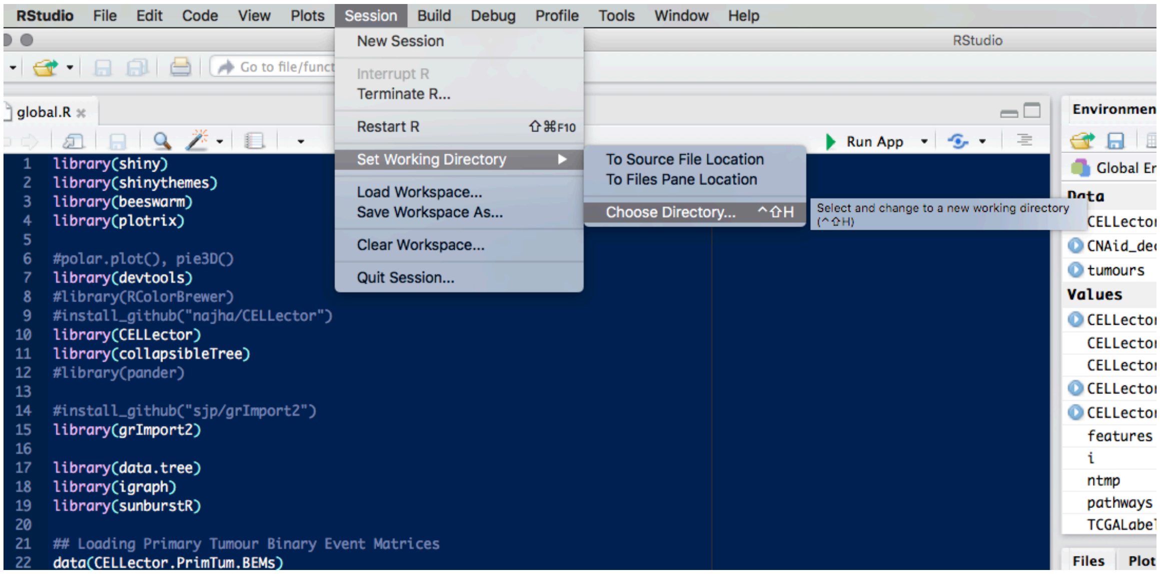

Finally, open the *global.R* script and click on *Run App*. All the required libraries should will be automatically installed or can be downloaded from (CRAN or Bioconductor) and installed manually.

Once the App is started, we strongly suggest to open it in your default web-browser to optimize the visualisation capabilities, by clicking on *Open in Browser* tab in Rstudio.

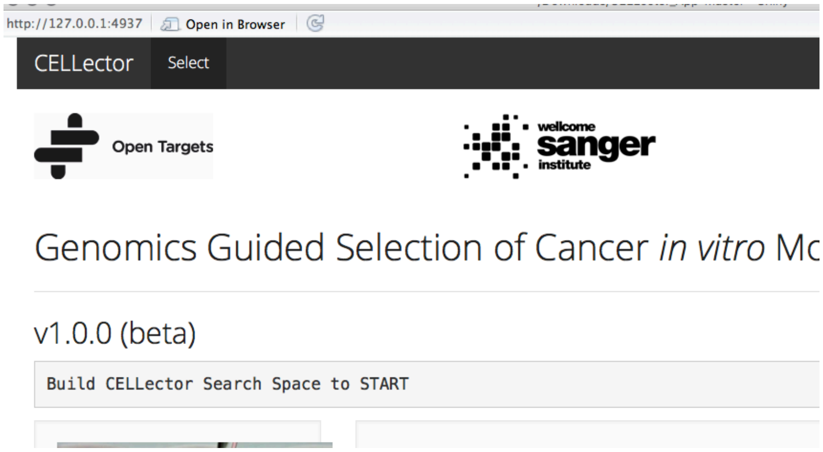
